# Genomic underpinnings of population persistence in Isle Royale moose

**DOI:** 10.1101/2022.04.15.488504

**Authors:** Christopher C. Kyriazis, Annabel C. Beichman, Kristin E. Brzeski, Sarah R. Hoy, Rolf O. Peterson, John A. Vucetich, Leah M. Vucetich, Kirk E. Lohmueller, Robert K. Wayne

## Abstract

Island ecosystems provide models to assess the impacts of isolation on population persistence. However, most studies of persistence have focused on a single species, without comparisons to other organisms they interact with in the ecosystem. The simple predator-prey system of moose and gray wolves on Isle Royale provides allows a direct contrast of genetic variation in a prey species with their natural predator. Wolves on Isle Royale exhibited signs of severe inbreeding depression, which nearly drove the population to extinction in 2019. In the relative absence of wolves, the moose population has thrived and exhibits no obvious signs of inbreeding depression despite being isolated for ∼120 years and having low genetic diversity. Here, we examine the genomic underpinnings of population persistence in the Isle Royale moose population. We document high levels of inbreeding in the population, roughly as high as the wolf population at the time of its decline. However, inbreeding in the moose population manifests in the form of intermediate-length runs of homozygosity indicative of gradual inbreeding, contrasting with the severe recent inbreeding observed in the wolf population. Using simulations, we demonstrate that this more gradual inbreeding in the moose population has resulted in an estimated 50% purging of the inbreeding load, helping to explain the continued persistence of the population. However, we also document notable increases in genetic load, which could eventually threaten population viability over the long term. Finally, we document low diversity in mainland North American moose populations due to a severe founder event occurring near the end of the Holocene. Overall, our results demonstrate a complex relationship between inbreeding, genetic diversity, and population viability that highlights the importance of maintaining isolated populations at moderate size to avert extinction from genetic factors.

**Significance statement:** Isolated wildlife populations face a high risk of extinction due in part to the deleterious consequences of inbreeding. Whether purifying natural selection can overcome these negative impacts by “purging” harmful recessive mutations is a topic of active debate. We characterized the extent of purging in an isolated moose population. Our results demonstrate signatures of gradual inbreeding in the population, ideal circumstances to facilitate purging. Using simulations, we demonstrate substantial potential for purging in the population, though we also show that fitness is reduced by small population size and inbreeding. Our findings provide insight into the mechanisms enabling persistence in isolated populations, with implications for conserving the growing number of isolated populations worldwide.

## Introduction

Anthropogenic habitat fragmentation has dramatically increased the number of isolated and inbred populations (1). To conserve these populations, a crucial question is whether they will be able to persist in isolation, or if they will be driven to extinction by deleterious genetic factors, such as inbreeding depression (2). Numerous examples exist of inbreeding depression driving population decline in isolated populations (reviewed in (3)). However, in some populations, harmful recessive mutations may potentially be ‘purged’ by purifying selection and such purging may avert inbreeding depression (2, 4–8). Purging may be most effective in populations where inbreeding is gradual due to a moderate population size (4–6, 9–11). However, the extent to which purging is a relevant factor for the conservation of threatened populations, and more broadly, the degree to which populations can persist with low genome-wide diversity, is controversial (9, 12–18).

One of the best-studied examples of inbreeding depression driving population decline is the gray wolf population on Isle Royale, an island in Lake Superior roughly 544 km^2^ in area. After ∼70 years of isolation at a population size of ∼25 individuals, the Isle Royale wolf population declined nearly to extinction, with just two individuals remaining in the population in 2018 (19). Recent research has demonstrated that this population collapse was a consequence of severe inbreeding depression in the form of widespread congenital deformities (20, 21). The decline of the Isle Royale wolf population allowed its main prey, moose, to thrive. The most recent moose census count was ∼2000 individuals, though the population generally numbers ∼1000 individuals (19). Moreover, despite the moose population having low genetic diversity and being isolated on the island for ∼120 years (22–25), it exhibits no obvious signs of inbreeding depression and has population growth rates similar to mainland populations (26). Thus, the contrasting fates of the Isle Royale wolf and moose populations provides a compelling case study for understanding the genetic underpinnings of population persistence in isolation and effects on predator-prey dynamics.

Outside of the Isle Royale population, North American moose are also known to have low genetic diversity relative to Eurasian moose, which is thought to be a consequence of a relatively recent founder event following the Last Glacial Maximum (27–29). Evidence for this recent founder also comes from a relative lack of population structure across North America as well as the near absence of moose in the North American fossil record prior to 15,000 years ago (27–30). Depending on how recent and severe this founding bottleneck was, the effects of purging associated with the bottleneck may still be apparent in the North American moose population. Thus, the ability of moose to persist in isolation on Isle Royale may be enhanced by purging from historical bottlenecks.

Here, we use a dataset of high coverage whole genome sequences from 20 North American moose and one Eurasian moose to characterize the impacts of bottlenecks, population isolation, and purging in North American moose, focusing on the Isle Royale population. We confirm previous findings of low genetic diversity in North American moose, especially Isle Royale moose, where levels of inbreeding are comparable to that of the Isle Royale gray wolf population at the time of its decline. Furthermore, we demonstrate that this low diversity is a consequence of severe founder events in both the North American and Isle Royale populations. Finally, we conduct extensive simulations exploring the impact of bottlenecks and population isolation on genetic load and purging in North American moose. These results suggest substantial purging associated with founding bottlenecks for the North American and Isle Royale populations. However, this purging also has been accompanied by a notable increase in genetic load. Overall, our analysis provides insight into how populations can persist despite severe bottlenecks and high inbreeding and emphasizes the importance of maintaining moderate population size to ensure viability in isolated populations. Moreover, our results highlight the differential impacts of inbreeding depression in isolated predator and prey populations, with implications for maintaining healthy ecosystems in the increasingly-fragmented landscape of the Anthropocene.

## Results

### Sampling and population structure

To examine patterns of moose genetic diversity in North America, we generated a high-coverage whole genome sequencing dataset for nine moose sampled from Minnesota and seven moose sampled from Isle Royale between 2005 and 2014. We added existing moose genomes to our dataset from Sweden, Alaska, Idaho, Wyoming, and Vermont. These genomes were aligned, genotyped, and annotated relative to the cattle reference genome (ARS-UCD1.2). Although a moose reference genome was recently published (30), we used the more distantly-related cattle reference in order to leverage its fully assembled chromosomes and high-quality annotations (see SI for further discussion). Average sequencing coverage after mapping was 21x (range 11-27; Table S1).

We first used these data to characterize population structure among North American moose, primarily aiming to assess evidence for isolation of the Isle Royale population. Principal component analysis (PCA) revealed a tight clustering of Isle Royale samples relative to other North American samples, which were distinctly clustered on the first PC (Fig. 1B). However, when down-sampled to one individual per North American population, the Isle Royale and Minnesota samples grouped more closely together, with overall patterns roughly reflecting North American geography (Fig. 1B, inset). Nevertheless, we observe notable differentiation between Isle Royale and Minnesota samples, with a mean F_ST_ = 0.083. These patterns were also reflected in a tree based on identity-by-state, which found a tight clustering of Isle Royale samples nested within other North American samples (Fig. 1C). Furthermore, using fastSTRUCTURE analysis we found no evidence for admixture between Isle Royale and mainland samples (Fig. 1D and S1-S2). Finally, we also estimated kinship for all North American samples, and found that the mainland samples are not closely related to one another (Fig. S3). However, two pairs of samples from Isle Royale exhibited kinship coefficients consistent with first-order relationships (mean kinship = 0.234; Fig. S3). In summary, these findings suggest that the Isle Royale population has been entirely isolated from nearby mainland moose populations as suggested by previous work (24, 25), and provide a general characterization of moose population structure in North America.

**Figure 1:**
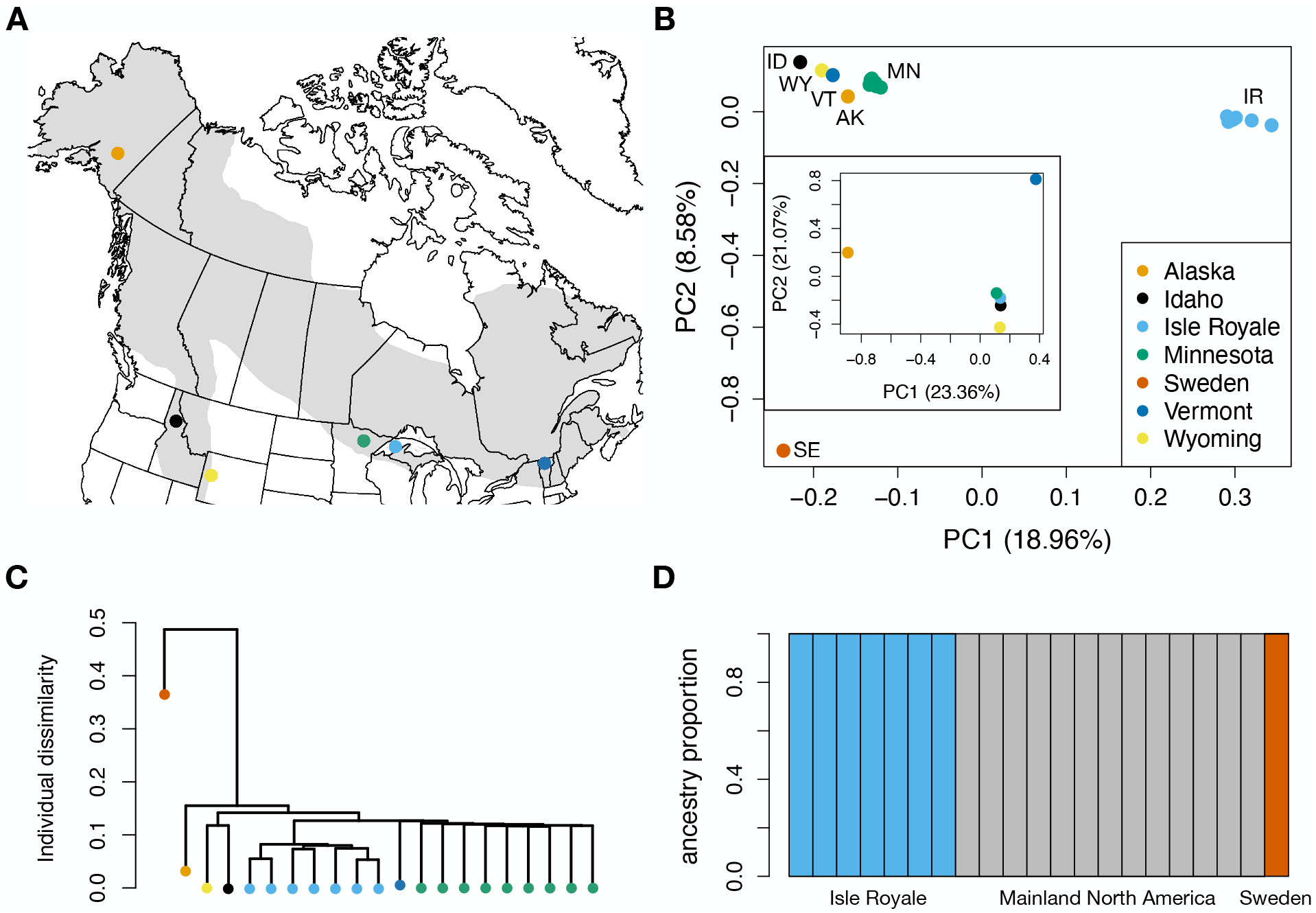
Moose sampling and population structure. (A) Map of North America including localities for individuals sampled for genomic data in our study. Note that Sweden is excluded. (B) PCA of 50,361 LD-pruned SNPs for all sequenced samples. Inset are results when down-sampling to one individual per population and excluding the Swedish sample. (C) Tree based on identity-by-state constructed using 50,361 LD-pruned SNPs. (D) fastSTRUCTURE results for K=3. See Fig. S1 for results with varying K values and Fig. S2 for results when down-sampling to four unrelated individuals each from Isle Royale and Minnesota.

### Genetic diversity and inbreeding

Next, we examined levels of genetic diversity and inbreeding across sampled individuals. Overall, we find that moose have relatively low diversity compared to other mammals (Fig. 2), though these estimates may be slightly downward biased due to using a distant reference genome (see SI for discussion). However, these biases do not impact estimates of relative diversity across moose populations, where several notable patterns are apparent. First, we observe substantially lower diversity in North American samples relative to a sample from Sweden, with a decrease of at least ∼34% (Fig. 2). This decrease in diversity is likely associated with a founder event for North American moose that is thought to have occurred during the last ∼15,000 years (27–29). We observe further reductions in diversity in the Isle Royale population, with an estimated reduction of ∼30% compared to samples from Minnesota (Fig. 2). Surprisingly, we find even lower diversity in mainland samples from Idaho, Wyoming, and Vermont, possibly due to these samples being near the southern range edge, where population densities are generally low and declining ((31); Fig. 2).

**Figure 2:**
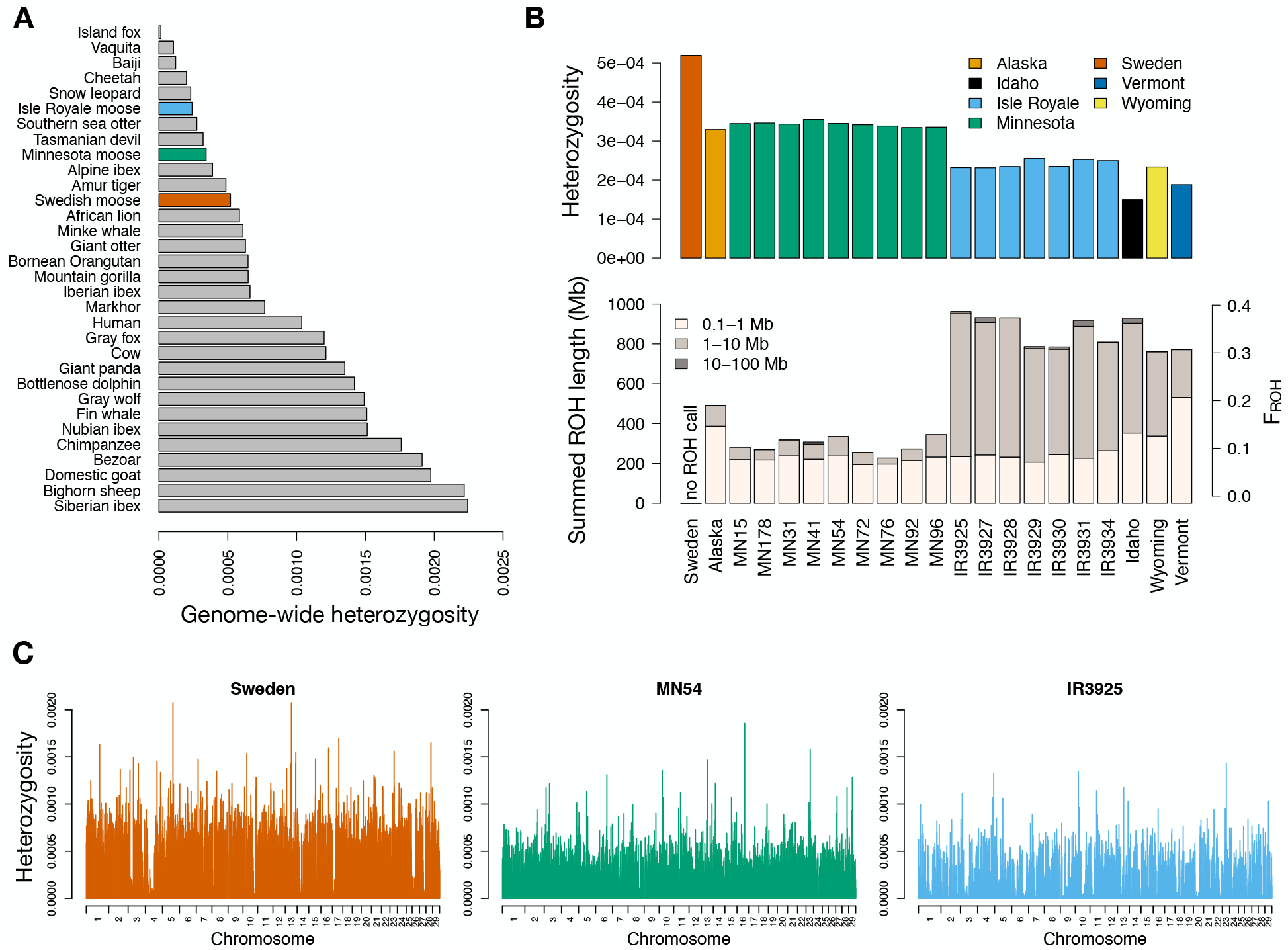
Moose genetic diversity and inbreeding. (A) Comparison of mean genome-wide diversity in three moose populations to published values for other mammals. (B). Plots of mean genome-wide diversity and summed ROH levels for North American moose genomes, with the corresponding F_ROH_ values on the right-hand axis. Note that we were not able to obtain ROH calls for the Sweden sample due to its differing population origin. (C) Per-site heterozygosity plotted in non-overlapping 1 Mb windows for representative individuals from Sweden, Minnesota, and Isle Royale. See Fig. S4 for plots of all individuals.

Mirroring these patterns of genetic diversity, the impact of inbreeding was prevalent across North American samples in the form of abundant runs of homozygosity (ROH), chromosomal segments that are inherited identical by descent from a recent common ancestor (32). Specifically, we observed high levels of inbreeding in samples from Isle Royale, Vermont, Idaho, and Wyoming, with ∼35% of their autosomal genomes being covered by ROH >100 kb on average (Fig. 2) and ∼26% covered by ROH >1 Mb (Fig. S5). As this fraction represents an estimate of the inbreeding coefficient (F_ROH_), this result suggests that these populations are on average more inbred than an offspring from a full-sib mating (F=0.25). Notably, these levels of inbreeding are comparable to the Isle Royale gray wolf population, where ∼20-50% of their autosomal genomes contained ROH >100 kb (20). By contrast, much lower levels of inbreeding were present in samples from Minnesota, Alaska, and Sweden, with ∼12% of these genomes covered by ROH >100 kb (Fig. 2), and ∼3% covered in ROH >1 Mb (Fig. S5).

### Demographic inference

To understand the demographic processes accounting for these patterns of genetic diversity and inbreeding, we fitted demographic models to the site frequency spectrum (SFS) using ∂a∂i (33). Briefly, this approach uses observed allele frequency information to estimate demographic parameters for a model with an arbitrary number of population size changes (epochs). Our first aim was to estimate the severity of the North American founding bottleneck, given the apparent impact of this bottleneck on observed levels of genetic diversity between Eurasian and North American moose (Fig. 2; (27)). We generated a folded SFS for our Minnesota sample, and inferred various population size change models including one, two, three, and four epoch models. Overall, the best-fitting model was a four-epoch model that included two ancestral epochs followed by a severe bottleneck to an effective population size (N_e_) of 49 for 29 generations and then expansion to N_e_=193,472 for the last 1,179 generations (Fig. 3). Bottlenecks that are mild with long duration can lead to similar patterns in the SFS as short and severe bottlenecks (34). Consequently, we found a similar fit for a model with a slightly more prolonged and less severe bottleneck of N_e_=218 for 142 generations followed by expansion to N_e_=105,531 for the last 1,223 generations (Table S2). Overall, both of these models are consistent in detecting a strong bottleneck of N_e_ = ∼50-225 for ∼30-150 generations followed by dramatic population growth taking place ∼1,200 years ago. The timing of expansion at ∼1,200 generations suggests a recent spread of moose across North America starting ∼9,600 years ago, assuming a generation time of 8 years (35).

**Figure 3:**
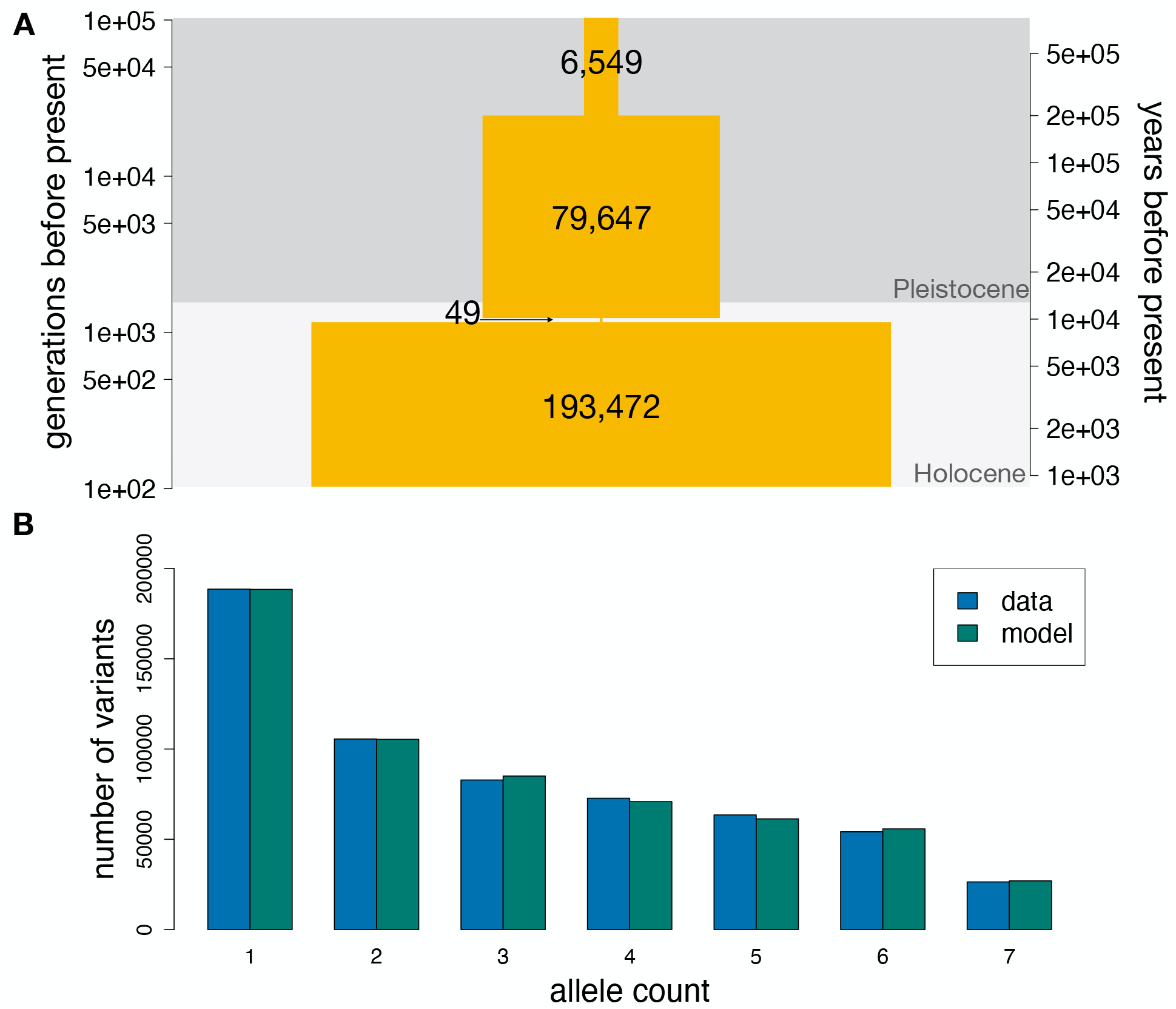
Demographic inference results. (A) Schematic of best-fit four epoch model based on the site frequency spectrum (SFS) for the Minnesota sample. Right-hand axis assumes a generation time of 8 years. Numbers denote maximum likelihood estimates of the effective population sizes at the various time points. Note the rapid and severe bottleneck occuring near the onset of the Holocene. See Table S2 for parameters of the second-best fitting run, which differs somewhat in bottleneck duration and magnitude and pre/post-bottleneck population sizes. (B) Comparison of the empirical projected folded SFS from the Minnesota sample with the SFS predicted by the model in shown in (A).

Our next aim for demographic inference was to obtain an estimate of the effective population size of the Isle Royale moose population after its founding ∼120 years ago using the SFS from our Isle Royale sample. Given the shared evolutionary history of the Minnesota and Isle Royale populations prior to their divergence, we fixed the demographic parameters of our four-epoch model inferred from the Minnesota samples (Fig. 3), then added a fifth epoch to this model representing the founding of Isle Royale. Furthermore, we fixed the timing of this fifth epoch to 15 generations ago, thus assuming that the population was founded in the early 1900s (120 years ago, assuming a generation time of 8 years; (35)), as suggested by available evidence (22, 23). We used this approach to retain power for estimating the Isle Royale effective population size when fitting a complex five-epoch model to an SFS from a small sample size. When fixing these parameters, we obtained an estimate of N_e_=187 on Isle Royale, highlighting a dramatic disparity in N_e_ between the North American and Isle Royale populations spanning three orders of magnitude. Additionally, given that the Isle Royale moose population on average numbers ∼1000 individuals (19), these results suggest an N_e_:N ratio of ∼0.19, consistent with those observed in other species (36). Notably, we observe the same N_e_:N ratio of ∼0.19 when comparing our estimated North American N_e_=193,472 (Fig. 3) to the current census estimate of one million (31).

### Quantifying putatively deleterious variation

To understand how the vastly reduced effective population size on Isle Royale may have impacted patterns of deleterious variation compared to mainland populations, we examined variants in protein-coding regions that were predicted to be putatively damaging or benign on the basis of evolutionary constraint (37). We observe a reduction in heterozygosity for both damaging and benign variants on Isle Royale, mirrored by an increase in homozygosity for the derived (i.e., mutant relative to the reference) allele (Fig. 4), as expected given the higher levels of inbreeding in the Isle Royale population. Specifically, we find that homozygous derived genotype counts are 9.7% higher for damaging variants and 6.8% higher for benign variants in Isle Royale moose compared to mainland moose. However, we do not observe an excess of derived alleles on Isle Royale (Fig. 4), as might be expected for a population that has accumulated an excess of weakly deleterious mutations due to relaxed purifying selection (38, 39). Collectively, these results suggest that the genetic load attributable to an accumulation of weakly deleterious mutations is negligible in Isle Royale moose.

**Figure 4:**
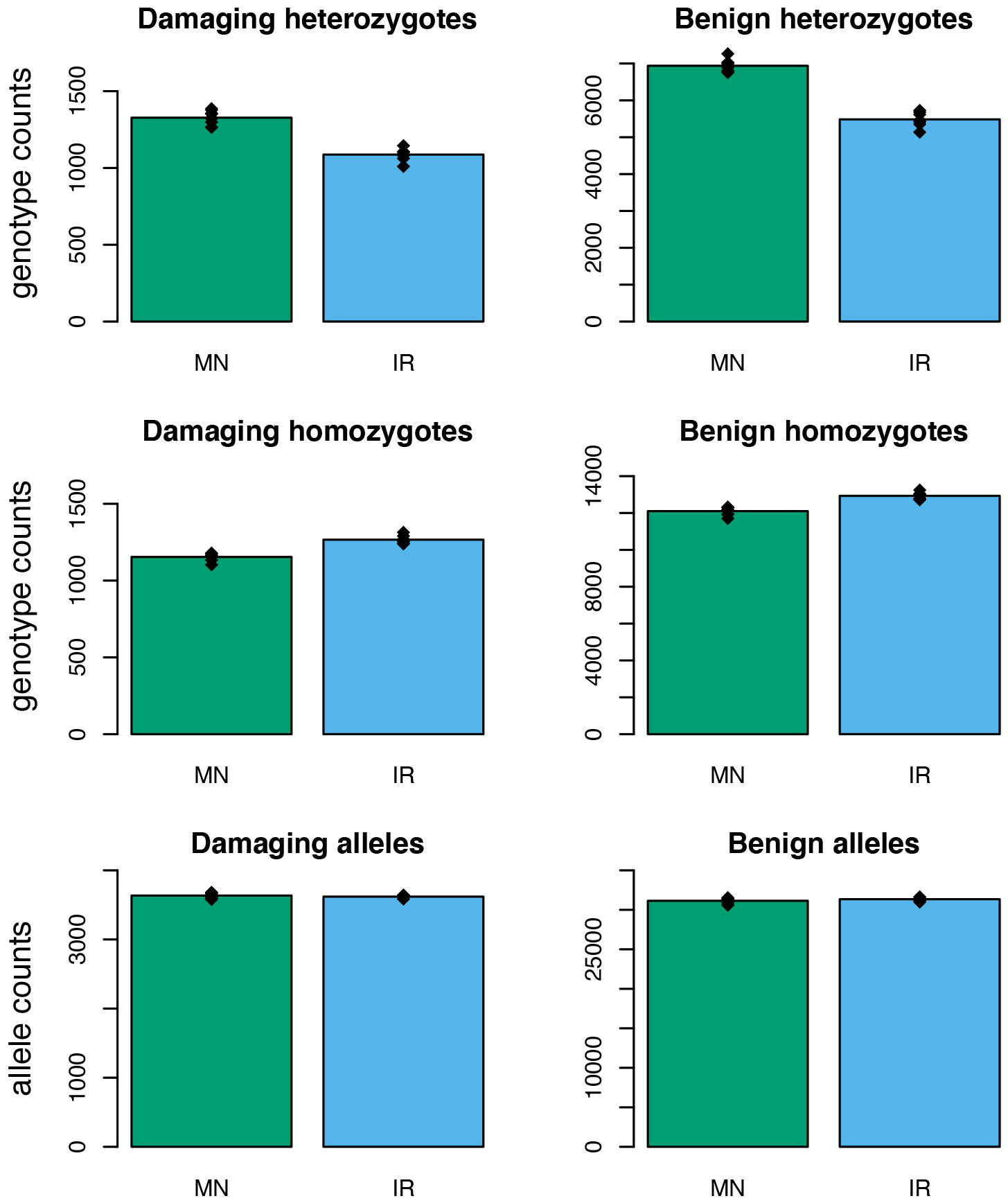
Empirical measures of deleterious variation in Isle Royale and Minnesota moose. Top row depicts counts of putatively damaging and benign heterozygotes, demonstrating that heterozygosity is reduced for both mutation types on Isle Royale. Middle row depicts counts of homozygotes for the derived allele at damaging and benign variants, similarly demonstrating increased homozygosity for both mutation types on Isle Royale. Bottom row depicts damaging and benign derived allele counts, demonstrating no differences between Isle Royale and Minnesota.

### Simulations of deleterious variation and genetic load

Empirical measures of deleterious variation are often challenging to interpret given that the functional impact and dominance of mutations are uncertain (40, 41). Consequently, we also conducted forward-in-time genetic simulations to assess the impact of bottlenecks on deleterious genetic variation in North American moose using SLiM3 (42). These simulations consisted of a 20 Mb chromosomal segment, which included a combination of introns, exons, and intergenic regions. Neutral and deleterious mutations occurred at a rate of 7e-9 per base pair (30), with deleterious mutations only occurring within exons. Selection coefficients for deleterious mutations were drawn from a distribution estimated from human genetic variation data (43), and dominance coefficients were assumed to be inversely related to selection coefficients, such that the most deleterious mutations were also the most recessive (see Materials and Methods).

Our first aim was to examine the impact of the North American colonization bottleneck on genetic diversity, genetic load, and purging. Here, we define “genetic load” as the realized reduction in fitness due to segregating and fixed deleterious mutations (44), and quantify purging as a reduction in the simulated “inbreeding load”, a measure of the quantity of recessive deleterious variation concealed in heterozygosis (2). To examine the dynamics of inbreeding, genetic diversity, and load in North American moose, we simulated under our best-fit demographic model (Fig. 3), which includes a founding bottleneck of N_e_=49 for 29 generations followed by expansion to N_e_=193,472 for 1,179 generations. Over the duration of this bottleneck, we observe a decrease in genetic diversity of 21%, along with a decrease in the inbreeding load of 24%, an increase in genetic load of 282% and an increase in F_ROH_ to 0.22 (Fig. 5). However, these increases in genetic load and F_ROH_ are largely absent after 1,179 generations of recovery, though levels of inbreeding notably remain above zero, in agreement with our empirical data (Fig. 2B). By contrast, genetic diversity and inbreeding load do not greatly increase after recovery, with the inbreeding load continuing to decline after the bottleneck and remaining 34% below its pre-bottleneck value even after 1,179 generations of recovery (Fig. 5). Thus, this result suggests that the North American moose population may still be experiencing the lingering purging effects of this founding bottleneck, despite occurring ∼9,600 years ago. Importantly, we observe qualitatively similar patterns when simulating under a model with a slightly longer and less severe bottleneck (Fig. S7), suggesting that these simulation results are robust to uncertainty in our estimated demographic parameters.

**Figure 5:**
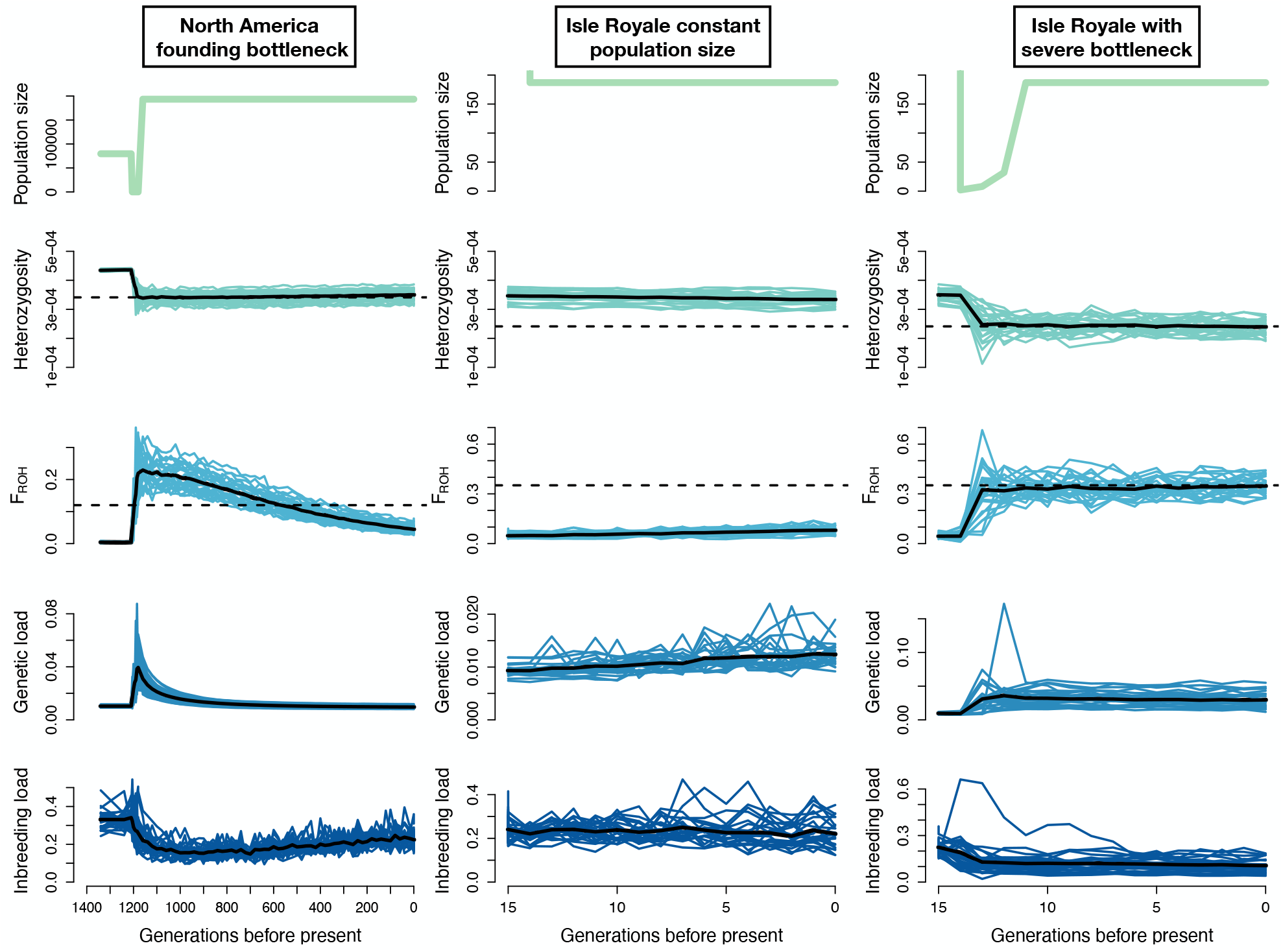
Simulation results under three demographic scenarios. Left column depicts simulation dynamics during the North America founding bottleneck; middle column depicts results when simulating the Isle Royale population at constant population size; right column depicts results when simulating the Isle Royale population including a severe founder event (N_e_={2,8,32} for the first three generations). Each column includes plots of the simulated effective population size, mean heterozygosity, mean levels of inbreeding (F_ROH>100kb_), mean genetic load, and mean inbreeding load from 25 simulation replicates. The black line represents the average from all replicates. The dashed lines represent the empirical estimates for heterozygosity and F_ROH_ from the Minnesota and Isle Royale populations, respectively. Note that the simulation trajectories do not reach these empirical estimates when assuming constant population size (middle column) but do when a founder event is included (right column). See Fig. S7 for results under additional bottleneck parameters.

Next, we examined the impact of isolation and small population size on Isle Royale on patterns of genetic variation and genetic load. We again simulated under our North America demographic model, though added a final epoch with the estimated Isle Royale demographic parameters of N_e_=187 for 15 generations. When simulating under this demography, however, we do not recapitulate the differences in genetic diversity and inbreeding observed in our empirical data between Isle Royale and mainland samples (Fig. 5). Specifically, heterozygosity decreased by only 3.6% compared to a ∼30% difference between Minnesota and Isle Royale samples in our empirical data, and levels of inbreeding increase only to F_ROH_=0.08 compared to F_ROH_=0.35 from our empirical data (Table S3).

We hypothesized that this discrepancy may be due to the absence of a severe founder event at the origination of the Isle Royale population in our model, given that the population is believed to be founded by a small number of individuals (22, 23). To test this hypothesis, we ran simulations where we included a bottleneck during the first three generations following the founding of Isle Royale. We tested three bottleneck severities with effective population sizes during the first three generations of N_e_={6,24,96}, N_e_={4,16,64}, and N_e_={2,8,32}, each followed by expansion to N_e_=187 for the final 12 generations. These bottleneck parameters were selected because available evidence suggests that population density was low soon after founding, particularly from 1900-1920, though it is unclear exactly how low or how many founders there were (22, 23). When varying these bottleneck parameters, we find that only the most severe bottleneck of N_e_={2,8,32} recapitulated the observed differences in genetic diversity and inbreeding, yielding a decrease in heterozygosity of 32% and increase in inbreeding to F_ROH_=0.35, in agreement with our empirical results (Figs. 5 and S7-S8; Table S3). Under this model, we also observe a relative increase in genetic load on Isle Royale of 206% as well as a 53% reduction in the inbreeding load (Fig. 5; Table S3). Thus, these results suggest that the Isle Royale moose population may have been founded by just two individuals, and that this severe founder event has been an essential factor in shaping patterns of genetic diversity, inbreeding, genetic load, and purging on the island. Finally, we do not observe any differences in allele counts between simulated island and mainland populations for mutations with selection coefficient (*s*) > -0.01 (Fig. S9), in agreement with our empirical result suggesting negligible impacts on load due to weakly deleterious mutations (Fig. 4). However, we do observe a sharp reduction in the number of strongly deleterious (*s* < -0.1) alleles per individual in the simulated Isle Royale population, suggesting that purging has largely been driven by a reduction in the number of strongly deleterious recessive alleles (Fig. S9).

Although our results suggest a substantial decrease in genetic diversity and increase in inbreeding in Isle Royale moose, field observations of the population have not detected obvious signs of inbreeding depression or reduced population growth rates (26). We hypothesized that this may be in part due to the purging that occurred during the North America founding event, which could enhance the ability of North American moose to persist at small population size. To test this hypothesis, we ran simulations under the above parameters including a severe Isle Royale founding bottleneck, but excluding the North America founding bottleneck. Here, we observe a much greater increase in genetic load on Isle Royale of 350%, compared to 206% when including the North America founding event (Table S3). Thus, these results suggest that the lingering effects of purging due to the North American founder event may have aided the ability of moose to persist at small population size on Isle Royale. In other words, the negative genetic consequences of small population size on Isle Royale may have been greater if the North American moose population had not experienced a strong bottleneck during colonization.

Next, we explored the potential impact of a low rate of historical migration on genetic variation in the Isle Royale population. Specifically, we explored the effect of a low rate of migration on genetic diversity, genetic load, levels of inbreeding, and inbreeding load. We ran simulations with migration fractions of 0.5% and 5%, roughly corresponding to 1 and 10 effective migrants per generation, respectively, chosen to model two relatively low but plausible rates of migration. Under the low migration scenario of 0.5%, results are nearly identical to the no migration scenario (Fig. S10; Table S3), implying that a very low level of historical migration (∼1 migrant per generation) would not have had much impact on the genetic state of the population. These results imply that we cannot fully rule out the possibility of a low rate of migration to Isle Royale, as suggested by direct observations of moose swimming between Isle Royale and the mainland (45). By contrast, when the migration fraction is increased to 5%, heterozygosity is higher and inbreeding lower relative to empirical values (Fig. S10; Table S3). In sum, these results further confirm that historical migration to Isle Royale was either absent or very low. Moreover, these results also suggest that any future attempts to restore genetic diversity and reduce genetic load in the Isle Royale moose population would require a relatively high rate of migration (>10 effective migrants per generation).

Finally, we explored the sensitivity of our results to selection and dominance parameters. Specifically, we simulated under parameters proposed by Kardos et al. (12), which assume that inbreeding depression is primarily due to recessive lethals and that deleterious mutations with *s* > -0.1 have largely additive effects on fitness. When simulating the North America founder event with these parameters, we observe a much smaller 22% increase in genetic load and a more substantial 60% decrease in the inbreeding load (Fig. S11). Additionally, the inbreeding load recovers much more rapidly following the bottleneck, due to the faster increase towards equilibrium of recessive lethal mutations (Fig. S11). When simulating a severe founder event for Isle Royale, we observe a much greater initial increase in genetic load; however, genetic load quickly decreases as recessive lethals are purged from the population, with a net increase of 66% (Fig. S12). Additionally, we observe substantial purging on Isle Royale, with a 75% reduction in the inbreeding load (Fig. S12). Thus, simulations under these parameters predict a much smaller increase in genetic load and much larger impacts of purging. This greater impact of purging is likely a consequence of the increased emphasis on recessive lethals in this model, which are most easily purged (5, 46).

## Discussion

Highly inbred populations are often thought to be doomed to extinction. However, some can persist, and understanding the factors enabling persistence can aid in conservation efforts. Our results document high inbreeding in the Isle Royale moose population (F_ROH_=0.35 on average; Fig. 2), roughly as high as the gray wolf population at the time of its decline. Yet, despite these high levels of inbreeding, the Isle Royale moose population does not exhibit obvious signs of inbreeding depression, and maintains population growth rates that do not noticeably differ from mainland moose (26). A key factor that likely underlies these different outcomes is the pace of inbreeding in these two populations: whereas the wolf population became quickly inbred while isolated at a population size of ∼25 for ∼70 years, inbreeding in the moose population was more gradual due to its more moderate population size of ∼1000 for a longer duration of ∼120 years. These differing demographic histories are reflected in the distribution of ROH lengths in the wolf and moose populations. In the wolf population, ROH were predominantly long (>10 Mb), reflecting recent and severe inbreeding (20), whereas the moose population exhibits an abundance of intermediate-length ROH (1-10 Mb; Fig. 2). Several recent studies have highlighted the severe fitness consequences of long ROH, which tend to be enriched for highly deleterious recessive alleles, whereas more intermediate-length ROH may be largely purged of such variation (20, 47–49). Although our results imply an elevated genetic load in the Isle Royale moose population (Fig. 5), this load has apparently not impacted population growth rates substantially, perhaps due to reduced interspecific competition on Isle Royale and soft selection (50). Overall, our results emphasize the importance of maintaining moderate size (N_e_ > 100) in isolated populations to enable purging and avert extinction in the short to intermediate term, in agreement with other studies (4–6, 9–11). Over the longer term, maintaining even larger population sizes (N_e_ > 1000) is preferable whenever possible to avoid the impacts of increasing drift load and loss of adaptive potential (12, 18).

Our results suggest that roughly half of the inbreeding load in Isle Royale moose may have been purged in the ∼15 generations or ∼120 years since founding (Fig. 5). The relatively rapid pace of this purging is notable, given that most existing examples of purging in wild populations occurred after thousands of years of isolation (4, 8, 51, 52). In Isle Royale moose, purging appears to have been accelerated by a severe founding bottleneck of perhaps just two individuals (Fig. 5). The impacts of severe bottlenecks on purging are well known (44), and have also been recently documented in an analysis of Alpine ibex genomes (7). For both Isle Royale moose and Alpine ibex, a severe bottleneck followed by relatively prompt recovery appears to have driven rapid purging on a timescale of ∼100 years. Thus, rapid purging on the timescale of anthropogenic fragmentation may only be possible in the presence of severe bottlenecks, perhaps precluding intentional purging <u>as</u> a viable conservation strategy. Nevertheless, many populations of at-risk species may have experienced historical purging due to severe bottlenecks or long-term moderate population size and identifying these populations could prove useful for future management actions.

Our findings also have important implications for understanding the evolutionary history and conservation status of mainland North American moose populations. Across all North American moose samples, we observe a reduction in genome-wide diversity of at least 34% relative to a sample from Sweden (Fig. 2), consistent with previous work (27, 30). Our demographic modeling indicates this reduction in diversity is due to a severe bottleneck in the ancestral North American moose population occurring ∼9,600 years ago (Fig. 3). This timing closely aligns with glacial recession at the onset of the Holocene 11,000 years ago as well as the North American fossil record (29). Furthermore, our simulation results suggest a substantial 34% purging of the inbreeding load associated with this founding bottleneck, the effects of which may persist until present day (Fig. 5). This phenomenon could further explain the success of the isolated Isle Royale moose population, implying that the founding individuals may have been ‘pre-purged’ of inbreeding depression. Moreover, the possibility of ‘pre-purging’ in North American moose could also help explain the success of other introduced moose populations in North American, such as the Newfoundland population, which was founded by just six individuals and now numbers >100,000 individuals (53). Nevertheless, many fragmented North American moose populations near the southern range edge have experienced recent declines (31). Though these declines have generally been linked to synergistic impacts of climate change and increasing disease and pathogen load (31, 54), the potential role of genetic factors has been largely overlooked. For example, we observed low genetic diversity in samples from Idaho and Wyoming (Fig. 2), perhaps due to the recent founding of these populations in the mid 19^th^ century and low population density (55). Notably, moose in this region exhibit low adult pregnancy rates (56), which could potentially be a consequence of inbreeding depression. Moreover, it is possible that low genetic diversity in these populations has increased their susceptibility to parasites (57). Overall, the causes of moose population declines near the southern range edge appear to be complex, and additional genomic sampling of these populations will be necessary to more fully investigate the potential role of genetic factors.

In conclusion, our results depict a complex relationship between genetic diversity, inbreeding, and population viability in isolated and fragmented populations. The contrasting fates of the Isle Royale wolf and moose populations serve as a dramatic example of the importance of maintaining isolated populations at moderate size to facilitate purging and avert extinction over the short to intermediate term. Moreover, this case study of predator and prey hints at a more far-reaching phenomenon, in which isolated predator populations may be doomed to extinction by inbreeding depression due to their naturally lower density, whereas the higher abundance of prey populations may enable them to purge the most severe impacts of inbreeding depression. In light of the well-documented connections among gray wolf, moose and plant abundance on Isle Royale (58), we suggest the possibility of an eco-evolutionary link between purging and the dynamics of the Isle Royale ecosystem. In general, purging may have system-wide effects in other isolated and fragmented ecosystems, where predator populations are declining in part due to inbreeding depression, and prey populations are thriving in their absence, often to the detriment of the broader ecosystem (59, 60). Thus, our results highlight a unique connection between deleterious genetic variation and ecosystem health, with implications for best management practices of small and fragmented populations.

## Materials and Methods

### Sampling and sequencing

Tissue samples were obtained opportunistically from moose carcasses on Isle Royale and Minnesota samples were collected during regular management activities by the Minnesota Department of Natural Resources (MN DNR). Isle Royale tissue samples were frozen and archived at Michigan Technological University and Minnesota tissue samples were provided by the MN DNR. DNA was extracted from samples using Qiagen kits and quantified using a Qubit fluorometer. Whole-genome sequencing was performed on an Illumina NovaSeq at the Vincent J. Coates Genomics Sequencing Laboratory at University of California, Berkeley and MedGenome. Existing genomes from (61) and (30) were downloaded from the National Center for Biotechnology Information (NCBI) Sequence Read Archive (see Table S1).

### Read processing and alignment

We processed raw reads using a pipeline adapted from the Genome Analysis Toolkit (GATK) (62) Best Practices Guide. We aligned paired-end 150bp raw sequence reads to the cattle genome (ARS-UCD1.2) using BWA-MEM (63), followed by removal of low-quality reads and PCR duplicates. Given that we do not have a database of know variants, we did not carry out Base Quality Score Recalibration, but instead carried out hard filtering of genotypes (see below). Although the cattle genome is highly divergent from moose, we opted to use it due to its much higher quality and contiguity compared to existing moose genomes (scaffold N50 of 103 Mb for ARS-UCD1.2 vs 1.7 Mb for NRM_Aalces_1_0) as well as its high-quality annotations and existing resources on the Ensembl Variant Effect Predictor database (64). To explore the potential impact of this on our downstream analyses, we also mapped a subset of nine genomes to the more closely related hog deer reference genome (ASM379854v1), which has high contiguity with a scaffold N50 of 20.7 Mb. Importantly, we found that the choice of reference genome here does not appear to qualitatively impact our genetic diversity and runs of homozygosity results. Thus, we use the cattle reference genome for all downstream analyses (see SI text for further discussion).

### Genotype calling and filtering

We performed joint genotype calling at all sites (including invariant sites) using GATK HaplotypeCaller. Genotypes were filtered to include only high-quality biallelic SNPs and monomorphic sites, removing sites with Phred score below 30 and depth exceeding the 99^th^ percentile of total depth across samples. In addition, we removed sites that failed slightly modified GATK hard filtering recommendations (QD < 4.0 || FS > 12.0 || MQ < 40.0 || MQRankSum < −12.5 || ReadPosRankSum < −8.0 || SOR > 3.0), as well as those with >25% of genotypes missing or >35% of genotypes heterozygous. We masked repetitive regions using a mask file downloaded from ftp://ftp.ncbi.nlm.nih.gov/genomes/Bos_taurus/. Finally, we applied a per-individual excess depth filter, removing genotypes exceeding the 99^th^ percentile of depth for each individual, as well as a minimum depth filter of six reads.

### Population structure and relatedness

We used SNPrelate v1.14 (65) to run principal component analysis (PCA), construct a tree based on identity-by-state (IBS), and estimate kinship among sampled genomes. For all analyses, we pruned SNPs for linkage (ld.threshold=0.2) and filtered out sites with minor allele frequency below 0.05, resulting in 50,361 SNPs for analysis. PCA was run both for all sampled individuals as well as for North American individuals down-sampled to one individual per population. We used the KING method of moments approach (66) to estimate kinship among North American moose samples. Finally, we estimated IBS among all samples, then performed hierarchical clustering on the resulting matrix to construct a dendrogram.

As another means of characterizing population structure, we used fastSTRUCTURE v1.0 (67) to test for admixture among sampled individuals. We converted our vcf to PLINK bed format with a minor allele frequency of 0.05 and maintained the order of alleles from the original vcf file. We ran fastSTRUCTURE on all sampled individuals as well as only Minnesota and Isle Royale individuals, each down-sampled to five unrelated individuals. For both analyses, we ran fastSTRUCTURE using values of *k* from 1-4. Finally, we used vcftools (68) to estimate Weir and Cockerham’s (69) FST between all Minnesota and Isle Royale samples using default settings.

### Genetic diversity and runs of homozygosity

We calculated heterozygosity for each individual in non-overlapping 1 Mb windows across the autosomal genome. We removed windows with fewer than 80% of sites called, as well as windows below the 5^th^ percentile of the total number of calls, as these windows have high variance in heterozygosity. We estimated mean genome-wide heterozygosity by averaging heterozygosity across windows for each individual.

Runs of homozygosity were called using BCFtools/RoH (70). We used the -G30 flag and allowed BCFtools to estimate allele frequencies. Due to the Swedish sample coming from a highly divergent population with differing allele frequencies, we excluded it from this analysis. We used a custom R script (71) to partition the resulting ROH calls into length categories 0.1-1 Mb, 1-10 Mb, and 10-100 Mb. We calculated FROH by summing the total length of all ROH calls >100 kb (or >1 Mb) and dividing by 2489.4 Mb, the autosomal genome length for the cattle reference genome. When conducting this analysis for the subset of samples mapped to the hog deer reference genome, we only used scaffolds >1 Mb in length, which together sum to 2479 Mb (∼93% of the total reference length).

### Identifying putatively deleterious variation

Variant sites were annotated using the Ensembl Variant Effect Predictor (VEP) v.97 (64). We used SIFT (37) to determine whether a nonsynonymous mutation is likely to be damaging or benign based on phylogenetic constraint. We classified protein-coding variants as “damaging” if they were determined to be “deleterious” nonsynonymous variants (SIFT score of <0.05) or variants that disrupted splice sites, start codons, or stop codons. Variants were classified as “benign” if they were determined to be “tolerated” nonsynonymous variants (SIFT score of ≥0.05) or synonymous mutations. Using these annotations, we tallied the number of derived alleles of each category relative to the cattle reference genome, as well as the number of heterozygous and homozygous derived genotypes, comparing these tallies for genomes sampled from Isle Royale and Minnesota. Variants that were that were fixed derived across the entire sample were ignored.

### Demographic inference

We estimated historical demographic parameters for North American moose based on the neutral site frequency spectrum (SFS) using ∂a∂i (33). In brief, we first focused on estimating parameters for the mainland North American population based on the neutral SFS for our nine Minnesota genomes, then used these results to guide inference of the effective population size on Isle Royale based on a neutral SFS from five genomes of unrelated Isle Royale individuals.

To generate a neutral SFS, we began by identifying regions that were >10kb from coding regions and did not overlap with repetitive regions (downloaded from ftp://ftp.ncbi.nlm.nih.gov/genomes/Bos_taurus/). We also excluded un-annotated highly conserved regions that are under strong evolutionary constraint, identified by aligning the remaining regions against the zebra fish genome using BLASTv2.7.1 (72) and removing any region which had a hit above a 1e-10 threshold.

We then generated a folded neutral SFS for these regions using a modified version of EasySFS (https://github.com/isaacovercast/easySFS), which implements ∂a∂i’s hypergeometric projection to account for missing genotypes. We found that the number of SNPs was maximized by using a projection value of seven diploids for the Minnesota sample and four diploids for the Isle Royale sample. In addition, we counted the number of monomorphic sites passing the projection threshold in neutral regions and added these to the 0 bin of the SFS.

We then used these SFSs to conduct demographic inference using the diffusion approximation approach implemented in ∂a∂i (33). Using the Minnesota SFS, we fit 1-epoch, 2-epoch, 3-epoch, and 4-epoch models. These models included the following parameters: Nanc (the ancestral effective population size), N1-3 (the effective size of the subsequent 1-3 epochs), and T1-3 (the duration of the subsequent 1-3 epochs; Table S2). In other words, a 3-epoch model includes the parameters Nanc, N1, N2, T1, and T2. Overall, we found the best fit for a 4-epoch model including expansion in the second epoch followed by a strong bottleneck and a final epoch of expansion, though with poor convergence of estimated parameters. Based on initial results, we constrained parameter space for the 4-epoch model by setting a limit on N1 to be in the range [10, 30]*Nanc, N2 to be in the range [1e-2, 5]*Nanc, and N3 to be in the range [10, 40]*Nanc.

We next sought to obtain an estimate of the effective population size on Isle Royale using a folded neutral SFS from five unrelated individuals, projected to four diploids. Given this limited sample size and the shared evolutionary history of Isle Royale and Minnesota moose, we fixed the parameters estimated from our 4-epoch model inferred above based on the Minnesota SFS. We then added a fifth epoch to the model, fixing the duration of this epoch to 15 generations, based on an estimated date of colonization of 1900 and 8 year generation time (35). Thus, the only estimated parameter in this approach is N5, the effective population size on Isle Royale.

We carried out inference by permuting the starting parameter values and conducting 50 runs for each model. We calculated the log-likelihood using ∂a∂i’s optimized parameter values comparing the expected and observed SFSs. For each model, we selected the maximum likelihood estimate from the 50 runs and used AIC to compare across models. We then used a mutation rate of 7e-9 mutations/site/generation and the total sequence length (L) to calculate the diploid ancestral effective population size as Nanc= Θ/(4*μ*L). We scaled other inferred population size parameters by Nanc and time parameters by 2*Nanc, in order to obtain values in units of diploids and numbers of generations.

### Simulations of deleterious genetic variation

We performed forward-in-time genetic simulations using SLiM v3.6 (42). We simulated a 20 Mb chromosomal segment with randomly generated introns, exons, and intergenic regions following the approach from (73). Thus, our aim with these simulations is not to quantify genome-wide effects of deleterious mutations, but rather to examine relative changes in deleterious mutations within a 20 Mb chromosomal segment. Deleterious (nonsynonymous) mutations occurred in exonic regions at a ratio of 2.31:1 to neutral (synonymous) mutations (74), and only neutral mutations occurred in intronic and intergenic regions. Following (30), we assumed a mutation rate of 7e-9 mutations per site per generation. Selection coefficients (*s*) for deleterious mutations were drawn from a distribution estimated using human genetic variation data by (43), consisting of a gamma distribution with mean *s* of -0.01314833 and shape = 0.186. Additionally, we augmented this distribution such that 0.5% of deleterious mutations were recessive lethal, given that this distribution may underestimate the fraction of lethal mutations (12). The dominance coefficients (*h*) of our simulations were set to model an inverse relationship between *h* and *s*, given that highly deleterious mutations also tend to be highly recessive (75, 76). Specifically, we assumed *h*=0.0 for very strongly deleterious mutations (*s* < - 0.1), *h*=0.01 for strongly deleterious mutations (−0.1 ≤*s* <-0.01), *h*=0.1 for moderately deleterious mutations (−0.01 ≤*s* <-0.001), and *h*=0.4 for weakly deleterious mutations (*s* > - 0.001). To test the sensitivity of our analysis to our assumed selection and dominance parameters, we also ran simulations under the selection and dominance parameters proposed by (12). Specifically, this model assumes that deleterious mutations come from a gamma distribution with mean *s* of -0.05 and shape = 0.5, augmented with an additional 5% of deleterious mutations being lethal. Dominance coefficients follow the relationship *h* = 0.5*exp(− 13\**s*); however, we simplified this to five dominance partitions for computational efficiency: *h*=0.48 for *s* ≥-0.01, *h*=0.31 for -0.1 ≤*s* <-0.01, *h*=0.07 for -0.4 ≤*s* <-0.1, *h*=0.001 for -1.0 ≤*s* <-0.4, and *h*=0.0 for *s*=-1.0. For all simulations, we retained fixed mutations, such that their impact on fitness was allowed to accumulate.

We set the population sizes of our simulations according to our best-fit 4-epoch demographic model based on the SFS from our Minnesota moose genomes (Fig. 3; Table S2). Specifically, this model estimated an ancestral effective population size of Nanc=6,548 diploids, followed by expansion to N1=79,647 for T1=22,628 generations, then contraction to N2=49 for T2=29 generations, and finally expansion to N3=193,472 for T3=1,179 generations. We also ran simulations under a second 4-epoch model that had similar log-likelihood and somewhat differing parameters of Nanc=7,017, N1=145,662, T1=20,883, N2=218, T2=142, N3=105,531 and T3=1,223. In both cases, we allowed the ancestral population to get to mutation-selection-drift equilibrium by running a burn-in at Nanc for 70,000 generations.

Following the fourth epoch of both models, we added a fifth and final epoch representing the founding of the Isle Royale population, consisting of Ne=187 for 15 generations. However, when simulating under this demography, we observed that the simulated levels of inbreeding and genetic diversity for the Isle Royale population did not recapitulate those observed in our empirical data (Fig. 5). Specifically, we observed only a 3.6% reduction in heterozygosity (compared to ∼30% in our empirical data) and an increase in FROH to just 0.08 (compared to 0.35 in our empirical data). We hypothesized that this was due to the lack of a founder event at the origination of the Isle Royale population in our model. To explore the impact of a founder event, we modified the effective population sizes during the first three generations of the Isle Royale population, using three plausible bottleneck parameters of Ne={6,24,96}, Ne={4,16,64}, and Ne={2,8,32}. We focused on the three initial generations after founding, reflecting the period from ∼1900-1924 when census estimates are crude and/or unavailable (22, 23). Specifically, little is known about the number of founding individuals, though it is likely this number was small, particularly if the population was naturally founded. Additionally, available records indicate a population size of ∼300 by 1920 and perhaps several thousand by 1930, suggesting that population growth was rapid following founding (22, 23). Following this three-generation bottleneck, we simulated the final 12 generations at our estimated Ne=187, representing an average effective population size for the period ∼1924-2020 when census estimates ranged from ∼500-2000 (average of ∼1000; (19)).

During simulations, we recorded mean heterozygosity, mean FROH for ROH >100 kb and >1 Mb, mean genetic load (calculated multiplicatively across sites), mean inbreeding load (measured as the number of diploid lethal equivalents), and the mean number of strongly deleterious (*s* < - 0.01), moderately deleterious (−0.01 ≤*s* <-0.001), and weakly deleterious (*s* > -0.001) alleles per individual. These quantities were estimated from a sample of 40 diploids every 1,000 generations during the burn-in, every 100 generations during the second epoch, every 5 generations during the North America founding bottleneck, every 20 generations during the fourth epoch, and every generation during the Isle Royale bottleneck. For all simulated scenarios, we ran 25 replicates.

## Acknowledgements

We are grateful to members of the Wayne and Lohmueller labs for helpful input on this work. We thank Michelle Carstensen and the Minnesota Department of Natural Resources for providing tissue samples used in this study. C.C.K. and K.E.L. were supported by National Institutes of Health grant R35GM119856 (to K.E.L.). A.C.B was supported by the Biological Mechanisms of Healthy Aging Training Program NIH T32AG066574. This work was supported by the National Science Foundation (DEB Small Grant #1556705).

## Data Availability

All scripts are available at https://github.com/ckyriazis/moose_WGS_project and raw data will be available on SRA upon publication.

## Author Contributions

C.C.K., R.K.W., and K.E.L. conceived the study. K.E.B., J.A.V., L.M.V., S.R.H. and R.O.P. acquired samples. C.C.K. conducted all analyses with input from A.C.B. and K.E.L. and wrote the manuscript with input from all authors. R.K.W. and K.E.L. jointly supervised this work.

