## Supplemental Information for "Genomic underpinnings of population persistence in Isle Royale moose"

**Supplemental Material**

**Impact of reference genome on estimates of heterozygosity and runs of homozygosity**

We explored the impact of reference genome selection on our analysis by mapping samples to two existing reference genomes: 1) the cattle genome (ARS-UCD1.2), which is very high quality with full chromosomes and a scaffold N50 of 103 Mb, but is highly divergent from moose (~26.3 My divergence according to timetree.org; (Kumar et al. 2017)), and 2) the hog deer genome (ASM379854v1), which is of somewhat lesser quality, lacking full chromosomes and with scaffold N50 of 20.7 Mb, but much less divergent from moose (13.5 My according to timetree.org). Although a moose reference genome exists (NRM_Aalces_1_0; (Dussex 2020) ; (Kumar et al. 2017)), it is highly fragmented (scaffold N50 of 1.7 Mb), posing issues for calling longer runs of homozygosity.

To guide reference genome selection, we initially mapped nine samples to both the cattle and hog deer reference genomes (Table S1). We observed much higher mapping rates to hog deer (98% on average, compared to 84% when mapping to cattle), as expected given the relative divergence of these species compared to moose. To explore the influence of these lower mapping rates and potential mapping bias on downstream analyses, we compared levels of heterozygosity and ROH calls for these nine samples following filtering. We found that heterozygosity was slightly lowered in samples mapped to cattle, though differences are small (3.7% on average; Fig. S12). However, relative patterns of diversity between moose samples appear not to be impacted, as we see a consistent pattern of reduced diversity in Isle Royale relative to Minnesota samples. Similarly, we observe slightly lowered $F_{ROH}$ estimates when mapping to cattle (5.7% lower on average; Fig. S13), though relative patterns between Isle Royale and Minnesota are again consistent. Additionally, reference genome appears not to impact our conclusion that ROHs in Isle Royale moose are predominantly of intermediate length (1-10Mb; Fig. S14).

Overall, we conclude that the selected reference genome does appear to have some influence on these results, though these impacts are relatively minor. Given this, we chose to use the cattle reference genome for our analysis to leverage the full chromosomes and high-quality annotations. More broadly, these results suggest that somewhat distant reference genomes may be a viable option in cases when high quality reference genomes do not exist for a given species.

**Table S1:** Summary of samples used in this study. Note that only nine samples were mapped to the hog deer reference genome for comparison with samples mapped to the cow genome. Also note that coverage was calculated for samples mapped to the cow reference genome.

| Individual | Source | Locality | Coverage | % reads mapped to cow | % reads mapped to hog deer |
|---|---|---|---|---|---|
| IR3925 | this study | Isle Royale | 27.6016 | 85.9 | 98.83 |
| IR3927 | this study | Isle Royale | 27.0643 | 82.71 | 98.16 |
| IR3928 | this study | Isle Royale | 25.5783 | 81.67 | 96.46 |
| IR3929 | this study | Isle Royale | 24.6561 | 86.47 | 98.33 |
| IR3930 | this study | Isle Royale | 19.9055 | 87.95 | - |
| IR3931 | this study | Isle Royale | 28.3286 | 87.19 | 98.98 |
| IR3934 | this study | Minnesota | 19.5286 | 87.68 | - |
| MN15 | this study | Minnesota | 21.9705 | 86.83 | - |
| MN178 | this study | Minnesota | 17.9505 | 85.88 | - |
| MN31 | this study | Minnesota | 24.3806 | 85.91 | 96.98 |
| MN41 | this study | Minnesota | 21.7691 | 82.51 | 97.96 |
| MN54 | this study | Minnesota | 25.2879 | 82.35 | 98.45 |
| MN72 | this study | Minnesota | 15.1187 | 84.86 | - |
| MN76 | this study | Minnesota | 17.8302 | 79.09 | - |
| MN92 | this study | Minnesota | 21.6431 | 88.23 | - |
| MN96 | this study | Minnesota | 22.5287 | 83.6 | 98.28 |
| C06 | Kalbfleisch et al. 2018 | Elk River, Idaho | 15.93829 | 75.36 | - |
| HM2013 | Kalbfleisch et al. 2018 | Paxon, Alaska | 16.05746 | 75.88 | - |
| JC2001 | Kalbfleisch et al. 2018 | Green River Lakes, Wyoming | 11.43739 | 78.14 | - |
| R199 | Kalbfleisch et al. 2018 | Lowell, Vermont | 19.26322 | 75.06 | - |
| Smoose | Dussex et al. 2020 | Gävleborg, Sweden | 20.10624 | 74.12 | - |

**Table S2:** Summary of ∂a∂i results based on the SFS from nine Minnesota samples projected to seven diploids. Note that all models showed convergence of parameter estimates and log-likelihoods, with the exception of the 4-epoch model, which had differing parameter estimates for the top two runs, though similar log-likelihoods. We therefore present parameter estimates and performed simulation analyses for both runs (see Figure S6 for comparison of simulation results). Note that K represents the number of estimated parameters and AIC = -2*LL+2*K.

| Model | K | Data log-likelihood | Log-likelihood | AIC | Parameter | Estimate |
|---|---|---|---|---|---|---|
| 1epoch | 0 | -45.61936776 | -1132.626666 | 2265.25 | Nanc | 12323 |
| 2epoch | 2 | -45.61936776 | -445.9806866 | 895.96 | Nanc | 12992 |
| | | | | | N1 | 9968 |
| | | | | | T1 | 2887 |
| 3epoch | 4 | -45.61936776 | -371.8761933 | 751.75 | Nanc | 13054 |
| | | | | | N1 | 92 |
| | | | | | T1 | 12 |
| | | | | | N2 | 3693967 |
| | | | | | T2 | 723 |
| 4epoch_model1 | 6 | -45.61936776 | -165.5233761 | 343.05 | Nanc | 6548 |
| | | | | | N1 | 79637 |
| | | | | | T1 | 22628 |
| | | | | | N2 | 49 |
| | | | | | T2 | 29 |
| | | | | | N3 | 193472 |
| | | | | | T3 | 1179 |
| 4epoch_model2 | 6 | -45.61936776 | -166.8463742 | 345.69 | Nanc | 7017 |
| | | | | | N1 | 145662 |
| | | | | | T1 | 20883 |
| | | | | | N2 | 218 |
| | | | | | T2 | 142 |
| | | | | | N3 | 105531 |
| | | | | | T3 | 1223 |

**Table S3: Summary of simulation results under various scenarios.** "IR bottleneck parameters" refers to the $N_e$ for each of the first three generations, followed by 12 generations at $N_e$=187 in each case. Note that we assumed a severe bottleneck ($N_e$={2,8,32}) unless otherwise noted. Percent changes and $F_{ROH}$ were all recorded at the end of 15 generations. Note that the empirical percent reduction in heterozygosity on Isle Royale is 29.6% and the empirical $F_{ROH}$ on Isle Royale is 0.35.

| simulated scenario | IR bottleneck parameters | % change in genetic load | % change in heterozygosity | % change in inbreeding load | $F_{ROH}$ |
|---|---|---|---|---|---|
| severe IR bottleneck | 2, 8, 32 | 206.1 | -31.6 | -52.6 | 0.347 |
| moderate IR bottleneck | 4, 16, 64 | 126.4 | -17.3 | -29.8 | 0.212 |
| weak IR bottleneck | 6, 24, 96 | 102.3 | -14 | -21.1 | 0.181 |
| no IR bottleneck | 187, 187, 187 | 32.7 | -3.6 | -3.6 | 0.081 |
| model2 parameters | 2, 8, 32 | 214.7 | -31.9 | 43.3 | 0.348 |
| no NA bottleneck | 2, 8, 32 | 350.1 | -32.4 | -38.4 | 0.326 |
| 0.005 migration fraction | 2, 8, 32 | 192.1 | -29.5 | -40.8 | 0.327 |
| 0.05 migration fraction | 2, 8, 32 | 65.5 | -8.8 | -22.9 | 0.133 |
| Kardos *hs* | 2, 8, 32 | 66.4 | -33.9 | -74.7 | 0.383 |

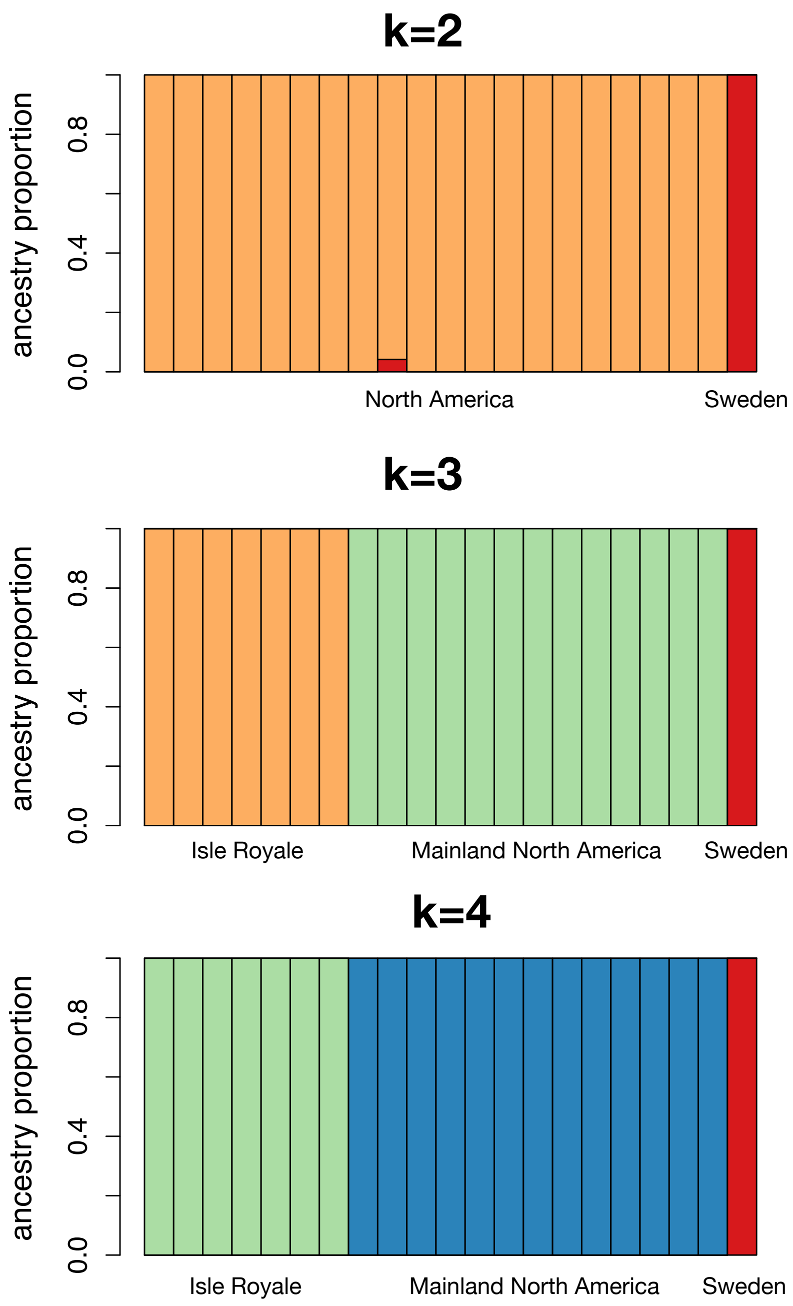


**Figure S1: Results for fastSTRUCTURE with different numbers of groups (k=2,3,4).** Colors indicate ancestry group. Results for k=3 are shown in Fig. 1. Note that no additional groups were detected when increasing from k=3 to k=4.

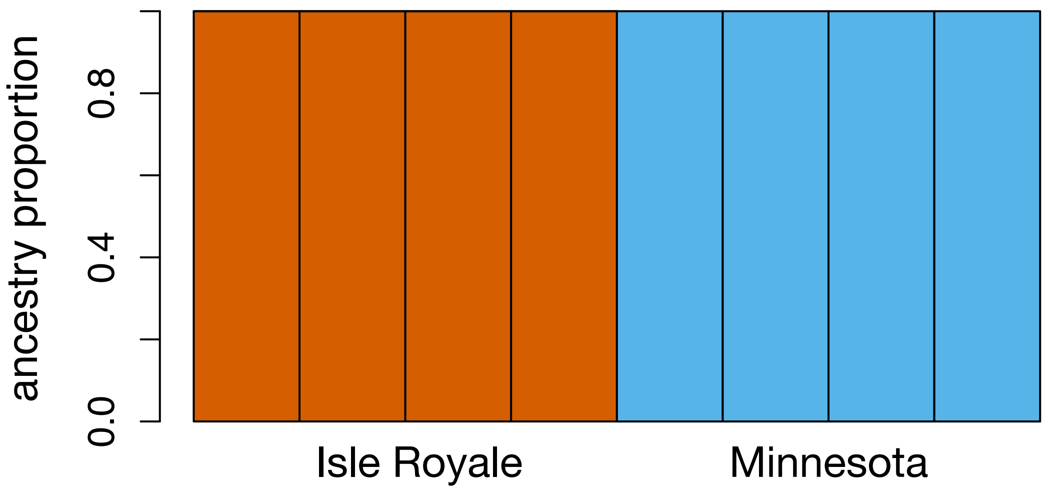


**Figure S2: fastSTRUCTURE results for K=2 when down-sampling to four unrelated Isle Royale and Minnesota individuals.**

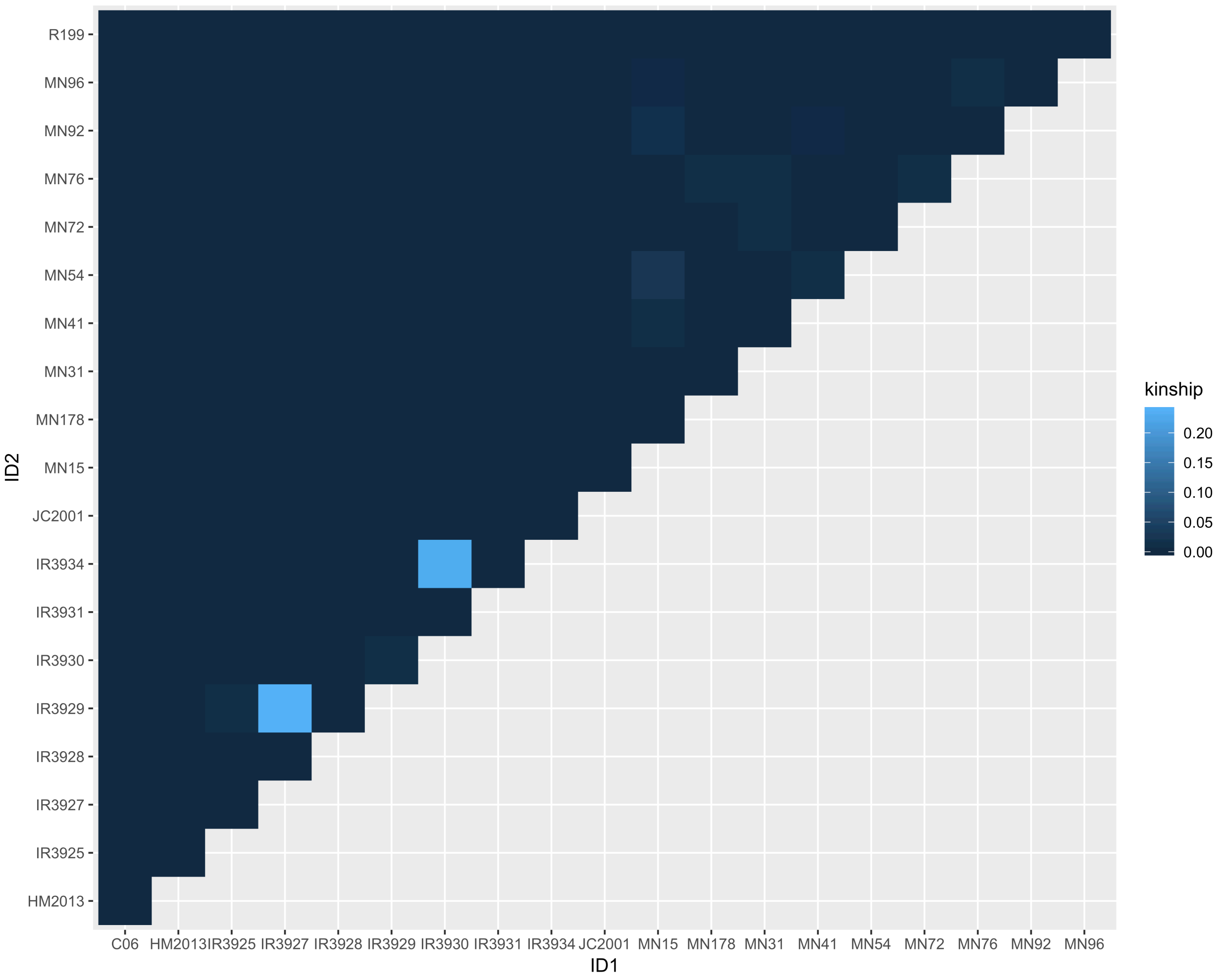


**Figure S3: Kinship coefficient estimates for North American moose samples.** See Table S1 for population information for each sample.

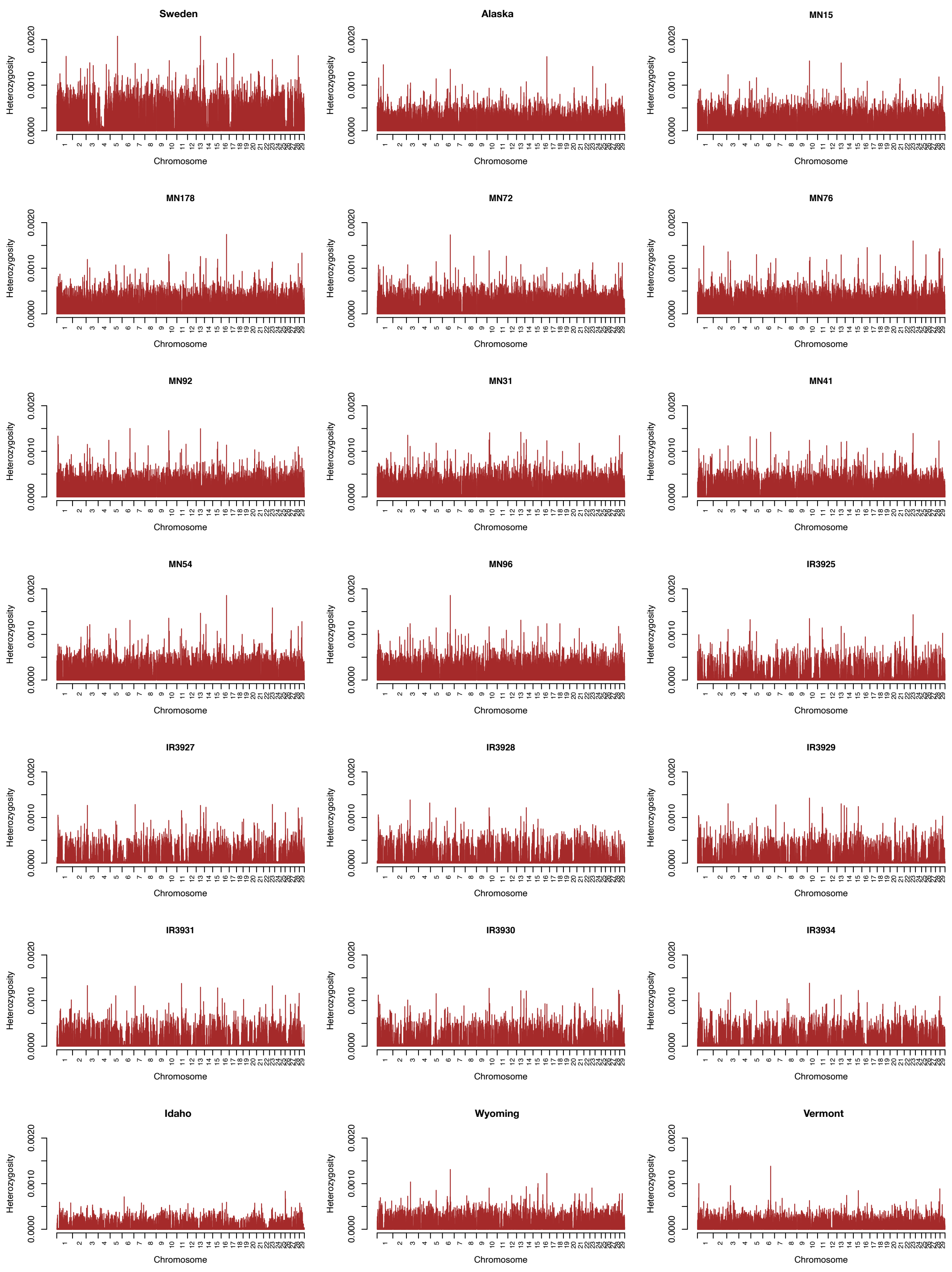


**Figure S4: Sliding window heterozygosity plots of all samples included in this study.**

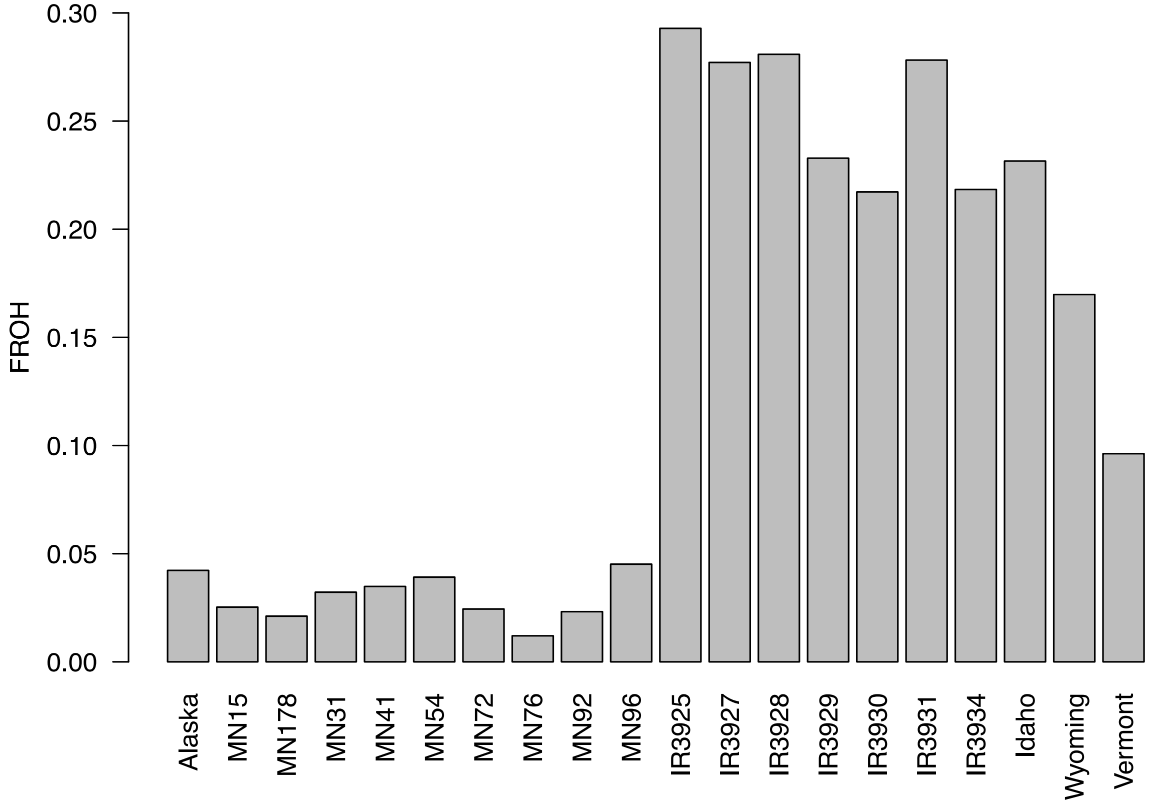


**Figure S5: Estimates of $F_{ROH}$ for all North American moose samples for ROH >1 Mb in length.** ROH were called using BCFtools/ROH.

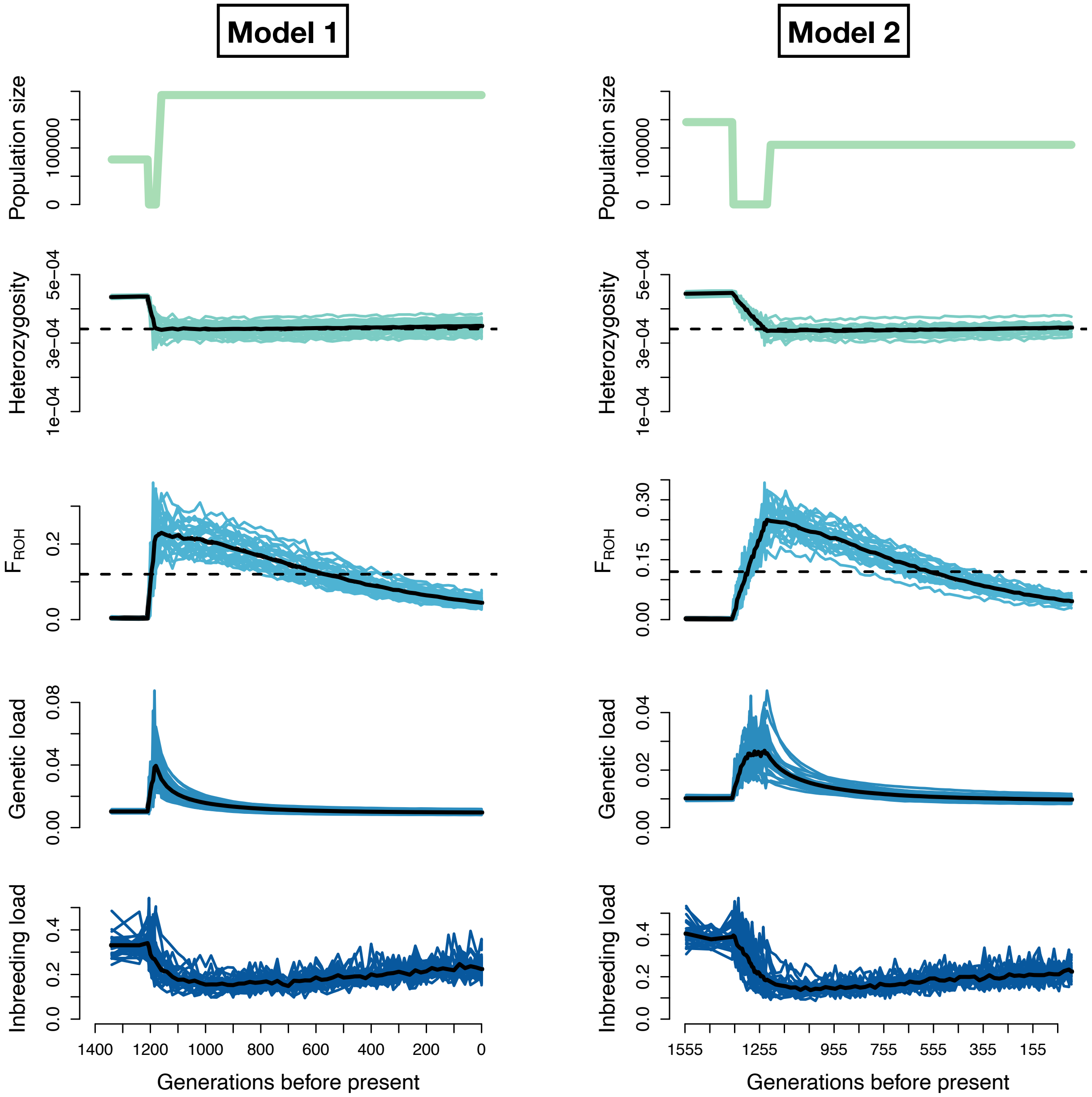


**Figure S6: Comparison of simulation results under demographic parameters from the top two ∂a∂i runs.** Note that the top two runs had similar log-likelihoods, though differed in estimates of several parameters (see Table S2 for details). However, these differences appear not to impact qualitative results. Note the differing y-axis scales.

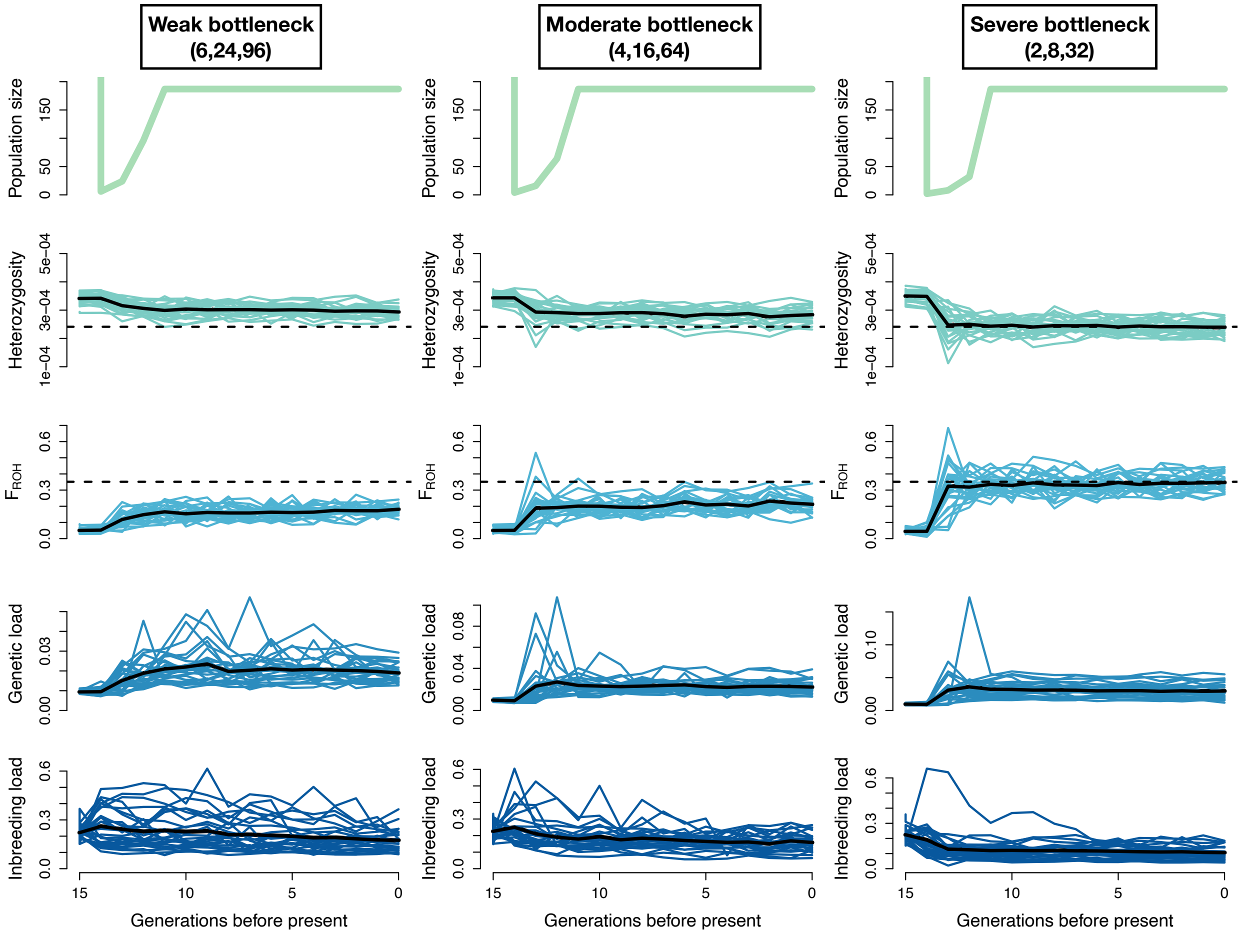


**Figure S7: Comparison of simulation results for the Isle Royale population under varying bottleneck strengths.** For each model, we assumed Ne=187 following a three-generation bottleneck of varying severity. Note that only the strongest bottleneck severity of Ne={2,8,32} recapitulates the observed heterozygosity and levels of inbreeding of the Isle Royale population (shown with dashed lines).

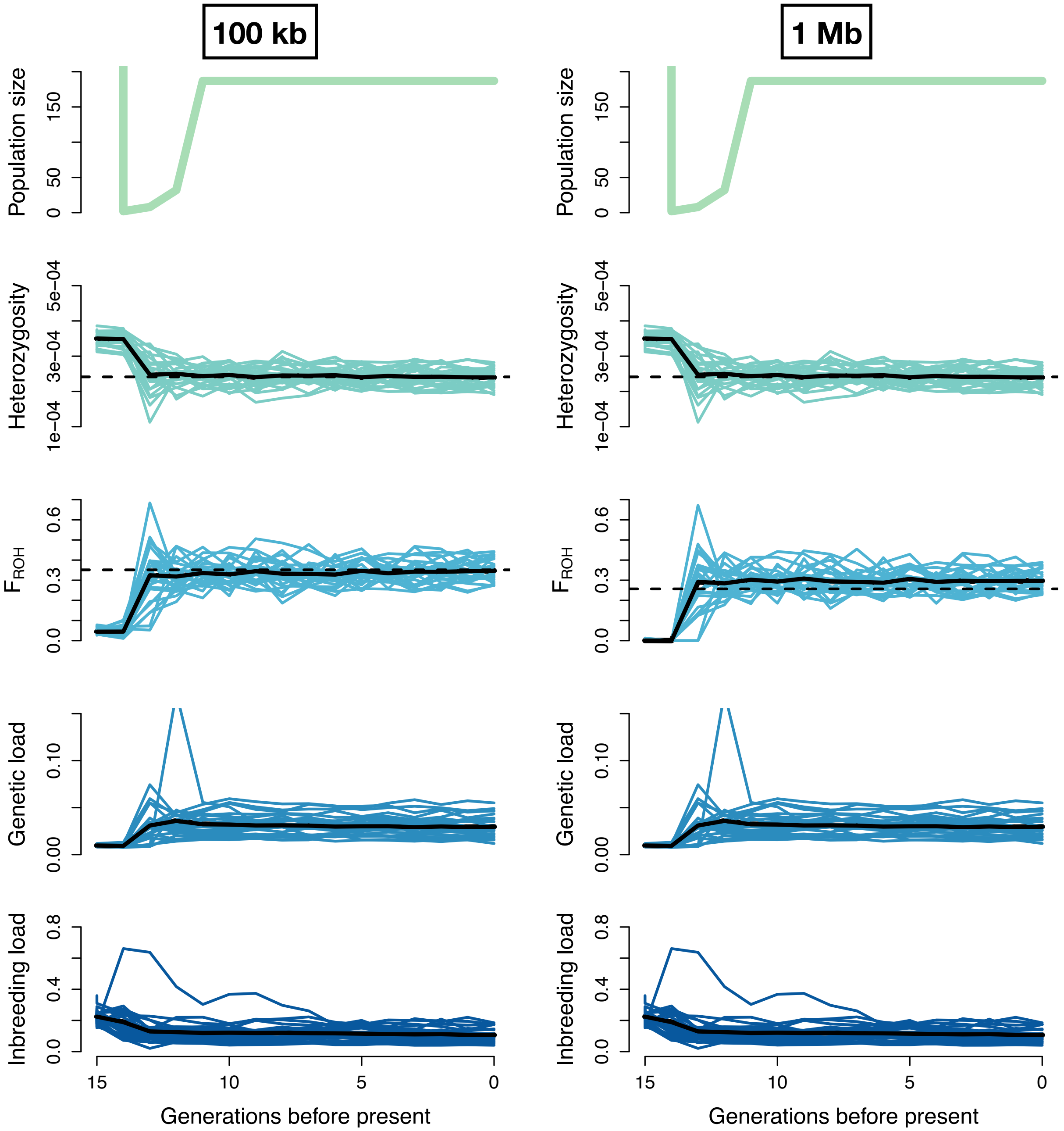


**Figure S8: Comparison of $F_{ROH}$ cutoff when comparing simulated bottleneck to empirical data.** The left panel depicts a simulated Isle Royale founding bottleneck when assuming an $F_{ROH}$ cutoff of 100 kb, both for the simulations and empirical data. The right panel depicts the same simulated bottleneck showing results with an $F_{ROH}$ cutoff of 1 Mb. Note that, in both cases, the simulated $F_{ROH}$ approaches a value close to the empirical $F_{ROH}$, suggesting that these simulated bottleneck parameters are in good agreement with the empirical ROH distributions on Isle Royale. In both cases, we assume a severe founding bottleneck with parameters $N_e$={2,8,32}.

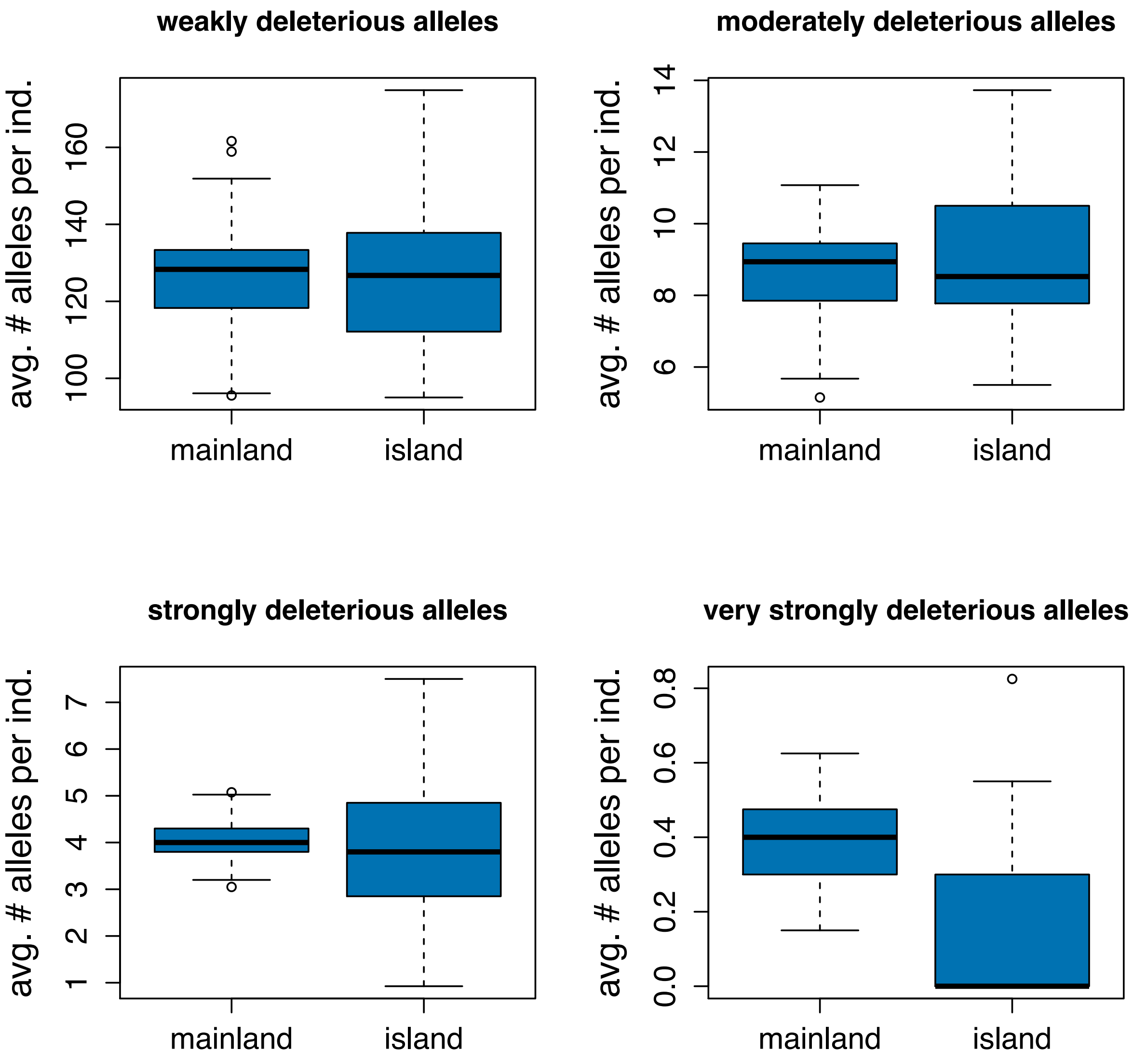


**Figure S9: Comparison of deleterious allele counts for simulated mainland and Isle Royale populations assuming a severe founder event scenario.** Allele counts for the Isle Royale population were recorded at the conclusion of 15 simulated generations. We define weakly deleterious alleles as mutations with $s > -0.001$, moderately deleterious alleles as $-0.01 \leq s < -0.001$, strongly deleterious alleles as $-0.1 \leq s < -0.01$, and very strongly deleterious alleles as $s < -0.1$. Note that minimal differences are apparent for all allele classes with the exception of very strongly deleterious alleles.

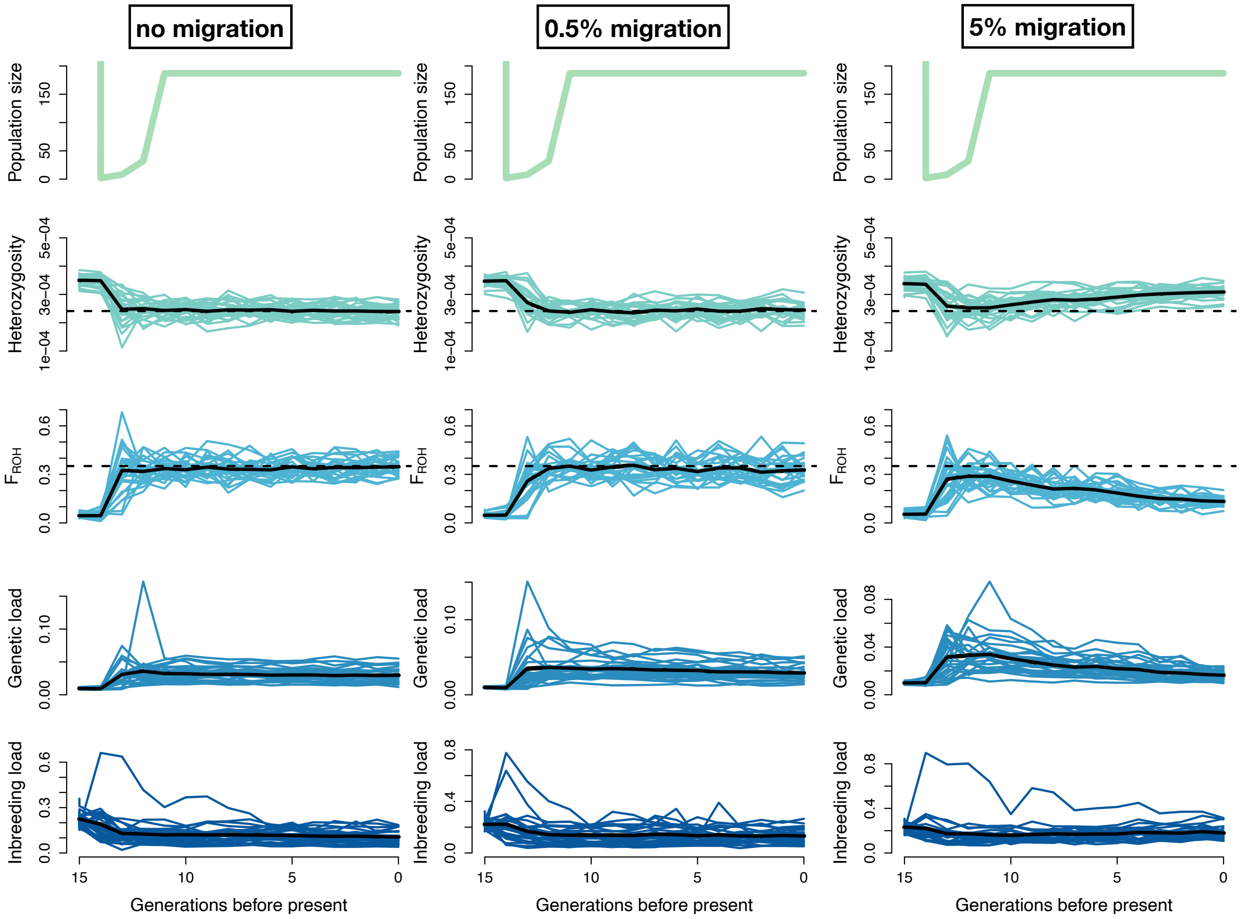


**Figure S10: Simulation results when including varying levels of migration to the Isle Royale population.** Left panel depicts scenario with no migration; middle panel shows results with a 0.5% migration fraction (roughly corresponding to one migrant per generation when $N_e$=187); right panel shows results with a 5% migration fraction (roughly corresponding to 10 migrants per generation). Note that the 0.5% migration results do not noticeably differ from the no-migration results, whereas the 5% migration results do. In all cases, we assume a severe founding bottleneck with parameters $N_e$={2,8,32}. Note the differing y-axis scales.

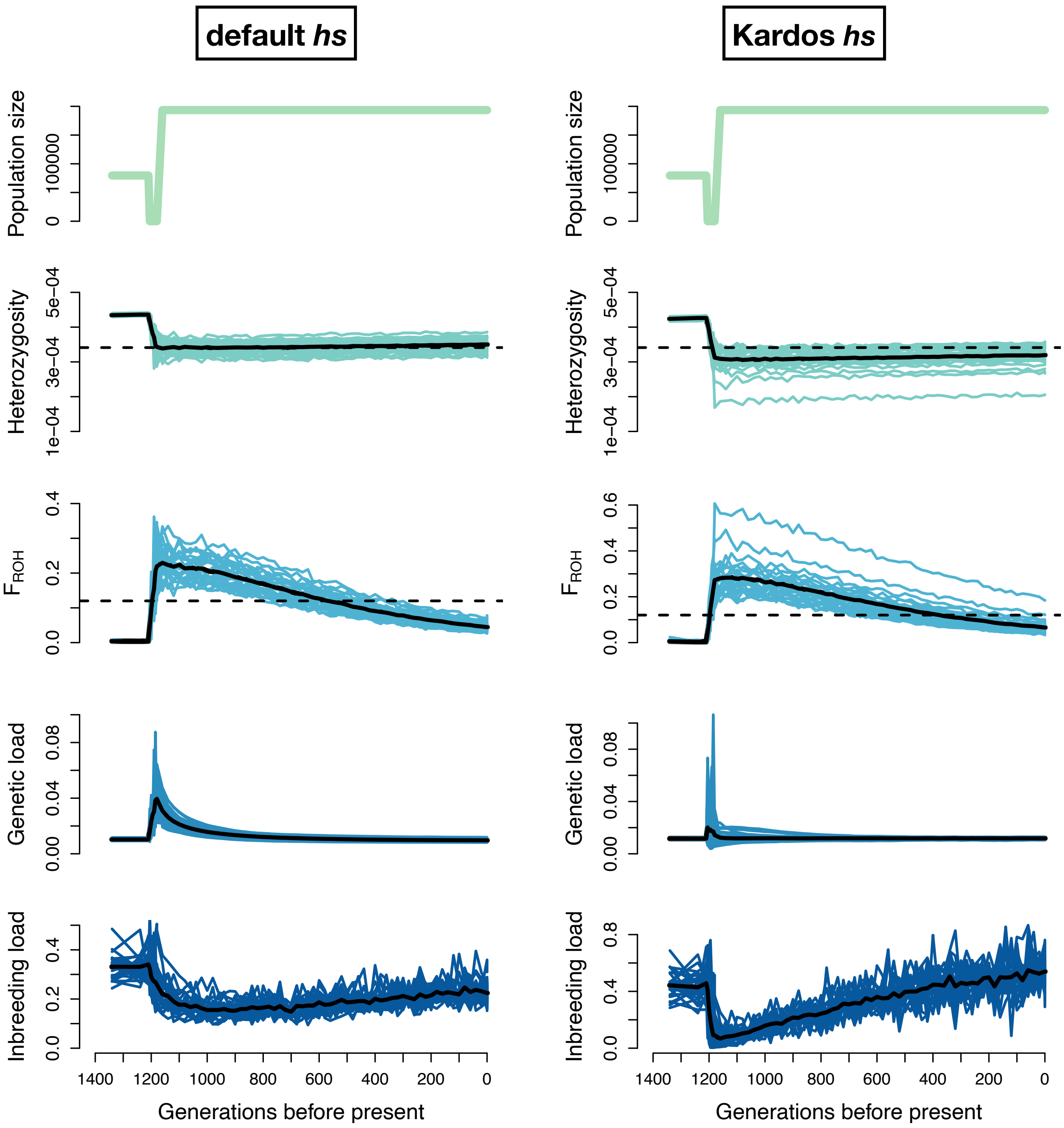


**Figure S11: Simulation results for the North America founder event under varying selection and dominance parameters.** Note the smaller increase in genetic load under the "Kardos" model during the bottleneck, and greater impact of purging. See Materials and Methods for details on model implementation. Note the differing y-axis scales.

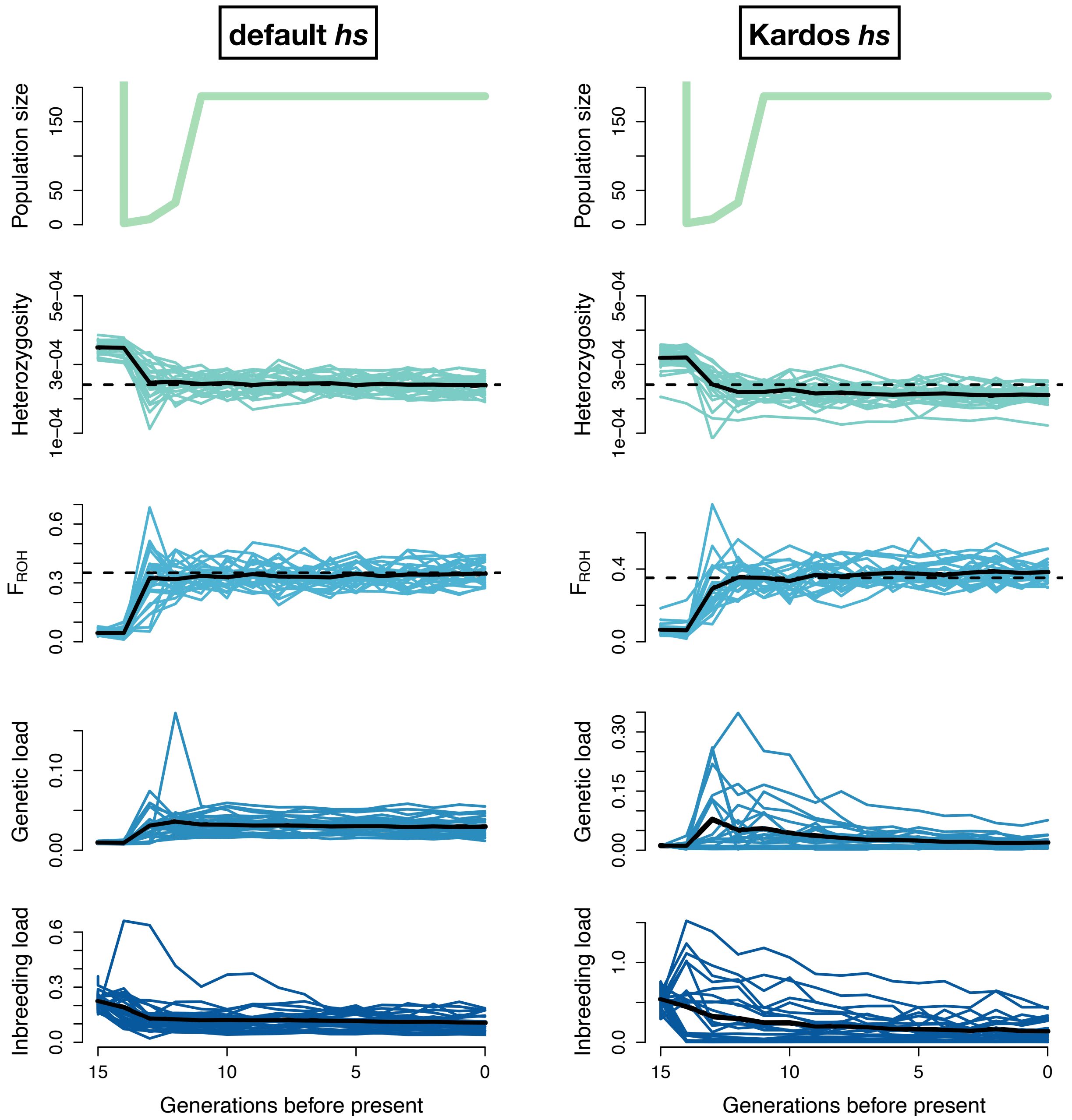


**Figure S12: Simulation results for the Isle Royale founder event under varying selection and dominance parameters.** In both cases, we assumed a severe founder event with $N_e$={2,8,32} for the first three generations. Note that, under the "Kardos" model, larger increases in genetic load occur initially but are later diminished due to greater purging. See Materials and Methods for details on model implementation. Note the differing y-axis scales.

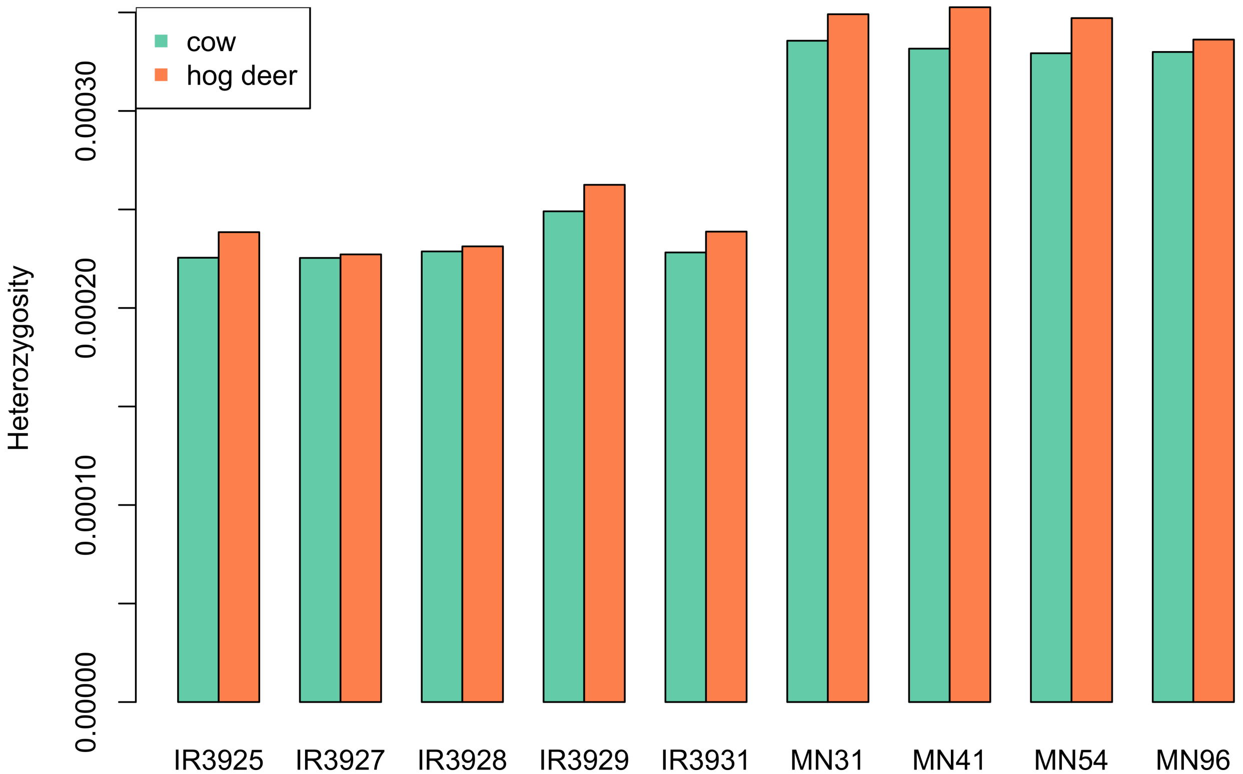


**Figure S13: Comparison of heterozygosity in a subsample of nine genomes aligned to the cow vs hog deer reference genome.** Note the slightly lowered heterozygosity when using the cow as a reference, due to the greater divergence between moose and cow.

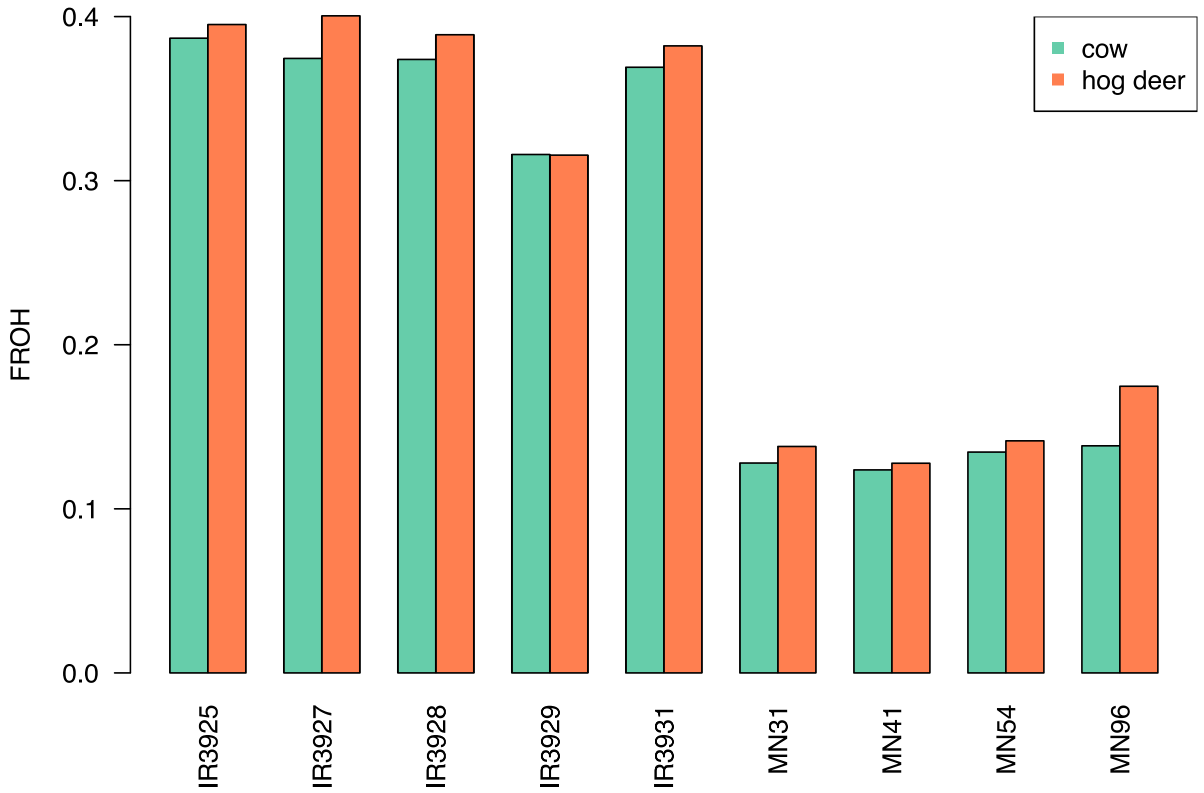


**Figure S14: Comparison of $F_{ROH}$ estimates in a subsample of nine genomes aligned to the cow vs hog deer reference genome.** For both cases, ROH calls were made using BCFtools using an ROH cutoff of 100kb. Note that estimates when mapping to cow are consistently lower, though estimates are overall in good agreement.

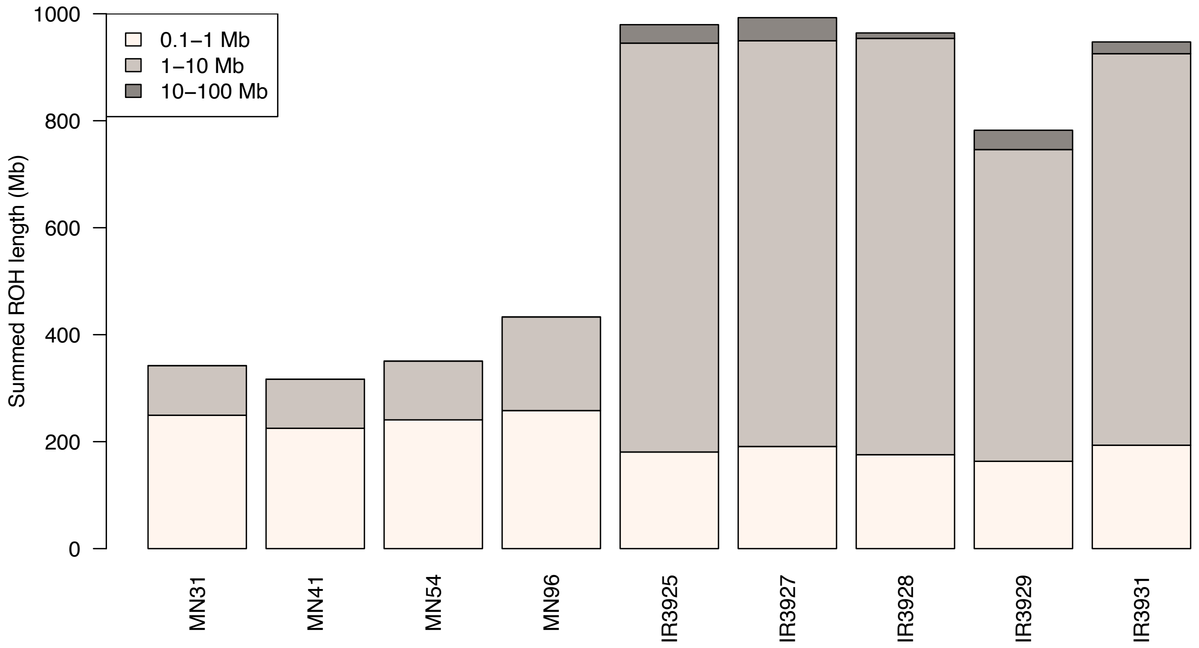


**Figure S15: Summed ROH lengths when mapping a subset of nine samples to the hog deer reference genome (ASM379854v1).** ROH were called using BCFtools. Note that ROH length distributions are highly similar to results using cow reference genome and BCFtools (see Fig. 2B), specifically with a large proportion of intermediate length (1-10Mb) ROH and very few long (10-100Mb) ROH in Isle Royale individuals.
